# Fixative eXchange (FX)-seq: Scalable Single-nucleus RNA Sequencing Analysis of PFA-fixed or FFPE Tissue

**DOI:** 10.64898/2026.03.05.709668

**Authors:** Han-Eol Park, Yu Tak Lee, Junsuk Lee, Hyelin Ji, Young-Lan Song, Jae Won Lee, Sung-Yon Kim, Junho K. Hur, Eunha Kim, Chung Whan Lee, Yoon Dae Han, Hyunki Kim, Chang Ho Sohn

## Abstract

Single-nucleus RNA sequencing (snRNA-seq) of clinical formalin-fixed, paraffin-embedded (FFPE) samples has long been a challenge due to low reverse transcription (RT) yields. Here, we present Fixative-eXchange (FX)-seq, a highly scalable snRNA-seq method for heavily paraformaldehyde (PFA)-fixed and/or FFPE samples. We employ an organocatalyst to facilitate the removal of PFA crosslinks to increase RT yield and additional regiospecific Pt(II)-based crosslinking of RNA molecules to prevent leakage. FX-seq reveals cellular heterogeneity across multiple fixed samples by analyzing 321,710 nuclei, including PFA-fixed tissue, FFPE blocks, thin FFPE and hematoxylin and eosin (H&E)-stained sections from mouse brain and human cancer specimens such as gastrointestinal stromal tumor and colorectal cancer. FX-seq enables integrated analysis with pathologist annotation to label tumor and non-tumor regions of H&E-stained sections. FX-seq can also be applied to PFA-perfusion-based animal studies, large human cohort studies, and personalized drug treatment through precision medicine.

## INTRODUCTION

In clinical settings, formalin-fixed, paraffin-embedded (FFPE) tissues serve as a primary resource for pathological diagnosis and long-term storage, and the number of their worldwide archives is estimated to exceed billions^1,2^. This may offer significant opportunities to obtain rich human transcriptome data at single-cell resolution, potentially enabling precision medicine to better understand and treat various human diseases^3,4^. Yet, single-cell or single-nucleus RNA sequencing of heavily fixed tissues such as FFPE tissues has been limited due to poor reverse transcription (RT) yields, even when using FFPE samples with reasonable RNA quality. Therefore, there is a strong unmet need for a high-quality, scalable single-nucleus RNA sequencing (snRNA-seq) method for heavily paraformaldehyde (PFA)-fixed and FFPE samples.

Recent studies have reported snPATHO-seq^5^, snFFPE-seq^6^, and snRandom-seq^7^, demonstrating the feasibility of snRNA-seq of FFPE tissue samples. However, snPATHO-seq employs pre-designed probe panels, which may bias the transcriptome landscape, while snRandom-seq uses random primers, which may result in inefficient mRNA data due to overwhelming reads from ribosomal RNAs (rRNAs). snFFPE-seq also showed limited scalability when analyzing human FFPE samples. Moreover, to improve the RT yield, these techniques rely on the conventional heat treatment at high temperatures and/or protease digestion of the tissue, which can lead to RNA degradation due to harsh heating or incomplete removal of the crosslinks due to the limited catalytic activity. In this regard, we envisioned that there is certainly much room for improvement by optimizing the process with a better understanding at the molecular level, which can further improve the data quality and scalability of FFPE snRNA-seq.

We began to speculate on the molecular mechanism of the inhibited RT process to improve *in situ* cDNA synthesis for extracted nuclei from heavily fixed samples. Previous studies have shown that PFA preferentially reacts with amino groups in nucleic acids to form crosslinks and dead-end methylol adducts^8–10^. Modifications of nucleobases decrease the RT yield for mRNA molecules due to the inhibition of Watson-Crick base pairing^8,11^. Moreover, a heavily modified poly(A) tail can reduce the efficiency of oligo(dT) primer annealing, and any hindrance to the progression of RTase can cause the enzyme to stall, leading to enzyme detachment and premature termination of cDNA synthesis^12,13^. Therefore, decrosslinking steps are critical to improve the quality of single-cell or snRNA-seq data from heavily fixed samples^14,15^ and also to prevent RNA degradation caused by harsh reaction conditions^16^.

Here, we report Fixative eXchange (FX)-seq to achieve high-quality snRNA-seq by incorporating chemical approaches with scalability. FX-seq enables (1) improved RNA retention by regioselective crosslinking, and (2) efficient reversal of PFA adducts using an organocatalyst under mild conditions (Figure 1A). Additionally, we employ combinatorial barcoding techniques for highly scalable nucleus labeling. However, frequent sample handling during nucleus isolation and combinatorial barcoding steps can lead to severe RNA degradation due to unwanted RNase contamination from unknown sources. Therefore, we further optimized the nucleus isolation process by employing the highly efficient, cost-effective chemical RNase inhibitor, polyvinyl sulfonic acid (PVSA), to enable scalable cell labeling.

**Figure 1.**
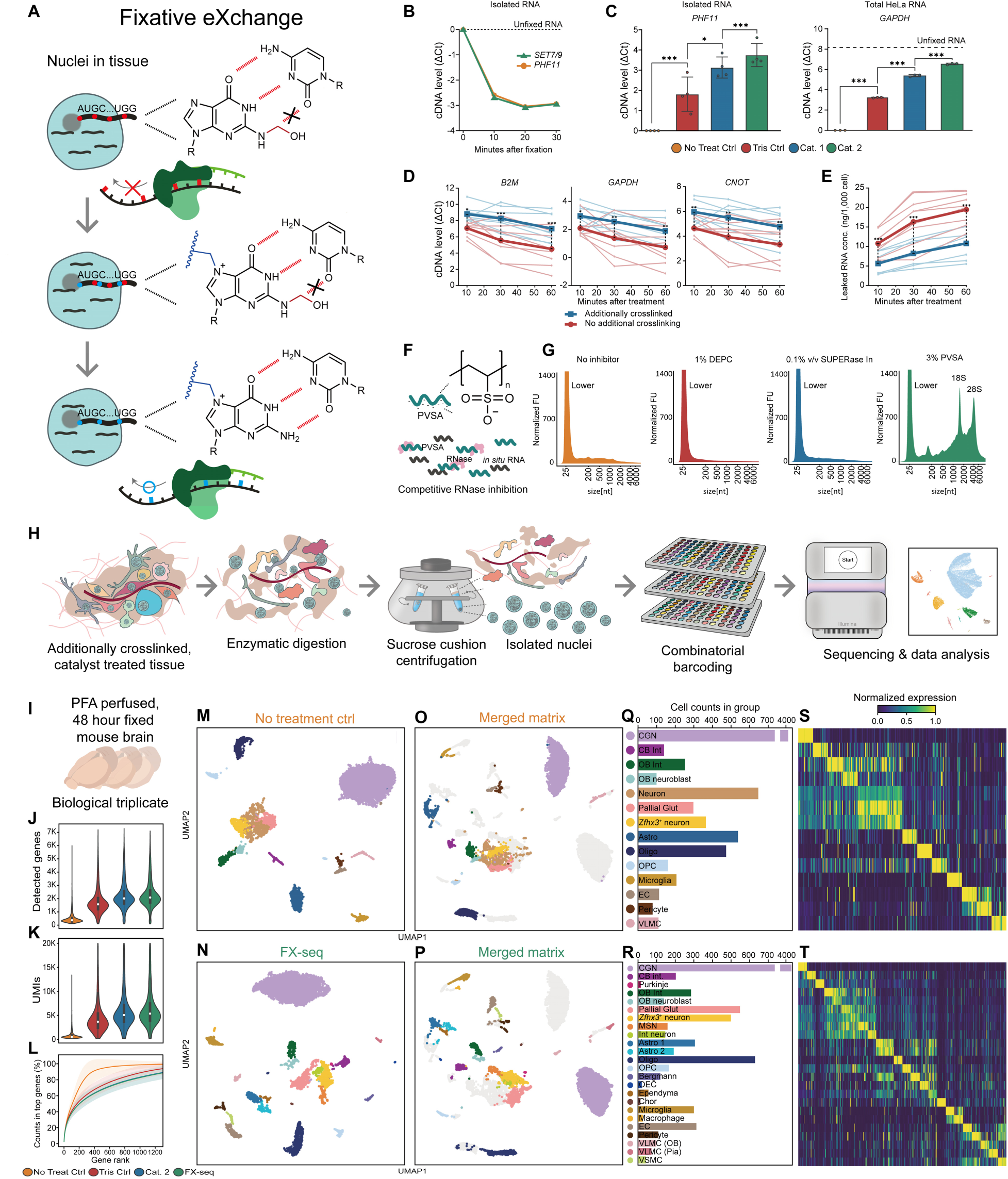
Concept of Fixative-eXchange (FX) and validation. (A) Schematic representation of the FX strategy. Heavy PFA fixation produces methylol adducts on primary amines that inhibit Watson-Crick base pairing and subsequent enzymatic activities. Additional crosslinking retains RNA molecules and an organocatalyst facilitates the removal of methylol adducts under mild conditions. (B and C) Relative cDNA synthesis yields were evaluated by normalization to the same amount of fresh RNA using qPCR measuring ΔCt values. PFA fixation inhibited the cDNA synthesis yield of isolated RNA (B). After the removal of PFA adducts with an organocatalyst, Cat. 2 shows better catalytic activities compared to Cat. 1 reported by Karmakar *et al.*^17^ (C). ΔCt values of each experimental group were normalized to the control group. Data are represented as mean ± SD. *P* values (NS ≥ 0.05, \**P* ≤ 0.05 \*\**P* ≤ 0.01 \*\*\**P* ≤ 0.001) were determined by unpaired one-way ANOVA followed by Bonferroni’s multiple comparison test. (D) Effects of additional crosslinking by regioselective crosslinkers measured in HeLa cells. Optimized conditions of Cat. 2 treatment improved cDNA yield but the prolonged reaction resulted in RNA leakage. The extent of RNA leakage during PFA removal was estimated by measuring ambient RNA levels per 1,000 cells by qubit and *in situ* cDNA yield by qPCR measuring ΔCt values. Data are represented as mean ± SD. *P* values (NS ≥ 0.05, \**P* ≤ 0.05 \*\**P* ≤ 0.01 \*\*\**P* ≤ 0.001) were determined by unpaired two-way ANOVA followed by Bonferroni’s multiple comparison test. (E) Additional crosslinking with our novel crosslinker minimized RNA leakage, resulting in improved RNA retention. Data are represented as mean ± SD. *P* values (NS ≥ 0.05, \**P* ≤ 0.05 \*\**P* ≤ 0.01 \*\*\**P* ≤ 0.001) were determined by unpaired one-way ANOVA followed by Bonferroni’s multiple comparison test. (F) PVSA as an effective RNase inhibitor for FX-seq. PVSA mimics negatively charged RNA phosphate backbones to inhibit RNase activity *in situ* by competitive binding. (G) Automated gel electrophoresis analysis of total RNA extracted from heavily PFA-fixed mouse brain tissue using various RNase inhibitors. PVSA shows the highest compatibility with the FX-seq protocol, which includes moderate heat treatment along with enzymatic digestion. Optimized nucleus isolation protocol with PVSA minimized RNA degradation during RNase-rich experimental procedures better than other inhibitors. Data are plotted by normalized fluorescence units (FU) per size (bp). “Lower” indicates low-molecular-weight species markers. (H) Optimized nuclei isolation strategy and the experimental sequences for FX-seq. Physically homogenized tissue is processed by FX treatment and enzymatically digested to isolate single nucleus suspensions and filtered by sucrose cushion centrifugation. Single nuclei are labeled by combinatorial barcoding using the sci-RNA-seq3 protocol with minor modifications. The sequencing library is then sequenced on an Illumina sequencer and the resulting sequencing data is analyzed to generate UMAP plots. (I–L) Littermate mouse brain triplicates heavily fixed with PFA were subjected to snRNA-seq (I) using four different conditions for PFA removal: isolated nuclei without any treatment (Ctrl), moderate heat treatment with 200 mM Tris buffer (Tris Ctrl), moderate heat treatment with Tris-buffered catalyst (Cat. 2), and the complete FX-seq procedure with additional crosslinking and tris buffered cat. 2 moderate heating (FX-seq). The individual components of FX-seq led to an increase in the number of detected genes (J), UMIs (K), and gene diversity (L). (M–T) UMAP clustering and cell-type annotation result of PFA-fixed control samples (n = 7497) and FX-seq (n = 8654) after doublet removal. Individual UMAP visualization and annotation (M and N) and UMAP analysis of merged gene expression matrix (O and P) demonstrated improved analytical resolution of cell type classification after FX-seq. Population distribution (Q and R) and specificity of marker genes is visualized (S and T). Abbreviations are as follows: CGN, cerebellar granule neuron; CB Int, cerebellar interneuron; OB Int, olfactory bulb interneuron; OB neuroblast, olfactory bulb neuroblast; Pallial Glut, pallial glutamatergic neuron; Astro, astrocyte; Oligo, oligodendrocyte; OPC, oligodendrocyte precursor; EC, endothelial cell; VLMC, vascular leptomeningeal cell; Purkinje, Purkinje cell; MSN, medium spiny neuron; Int neuron, interneuron; Bergmann, Bergmann glia; OEC, olfactory ensheathing cell; Ependyna, ependymal cell; Chor, choroid plexus epithelial cell; VLMC (OB), vascular leptomeningeal cell in olfactory bulb; VLMC (Pia), vascular leptomeningeal cell in pia mater; VSMC, vascular smooth muscle cell.

## RESULTS

### Optimization of FX and nucleus isolation

To address the aforementioned challenges associated with PFA fixation, we first adopted an approach by Karmakar *et al.*^17^ using organocatalysts to promote the removal of PFA adducts under mild conditions. Before testing catalysts, we confirmed that RT inhibition does indeed occur through PFA crosslinking (Figure 1B). We then further screened catalyst candidates for better catalytic activities and found that cat. 2 outperformed the previously reported catalyst with the best reaction kinetics observed for fixed isolated RNA species and total HeLa RNA (Figure 1C and S1A and S1B)^18^.

Next, we proceeded to optimize protocols for fixed HeLa cell cultures (Figure 1D). To maximize RT efficiency, we hypothesized that the extended catalytic treatment could further improve the *in situ* RT yield. However, we found that the non-cytosolic, ambient RNA concentration increased, indicating leakage from the fixed cells (Figure 1E). To avoid this loss, we further crosslinked the cells with a regiospecific Pt-based chemical crosslinker, synthesized by coupling guanine-N7-specific cisplatin derivatives with a di-amino-undeca-polyethylene glycol (PEG) linker (Figure S2A–S2C; STAR Methods). This crosslinker can provide additional support for RNAs by forming bonds at guanine-N7 residues in a regioselective manner (Figure S2D) without interfering with the Watson-Crick base pairing^19,20^. Using isolated RNA species, we confirmed that regiospecific crosslinking at guanine-N7, without disruption of Watson-Crick base pairing, has a minimal effect on RTase processivity and therefore maintains cDNA yield compared to unfixed RNA (Figure S3A–S3J). We then balanced the degree of additional *in situ* crosslinking to minimize RNA leakage without compromising the yield of PFA adduct removal and ultimately improve cDNA yield (Figure 1D and E; STAR Methods).

To extend the applicability of FX-seq to more complex biological contexts, we further optimized the nucleus isolation procedure for heavily fixed tissues and organs. We found that the optimized protocol allows robust extraction of nuclei from various mouse organs (Figure S4 and S5; STAR Methods). However, during the sample manipulation associated with nucleus isolation and heating for PFA removal, RNA quality was significantly compromised by unknown RNase activities or RNA autolysis (Figure 1G). To address this issue, we searched for an effective and cost-effective RNase inhibitor (RI) to replace expensive protein-based RIs. We identified PVSA, which is thermostable and compatible with the downstream enzymatic reactions. Unlike protein-based RIs, PVSA inhibits RNase activity by blocking the active site as a non-reactive competitor, leaving RNAs nascent without RI adducts, potentially beneficial for improving the *in situ* RT yield^21^. Under the moderate heating condition for the catalytic treatment, PVSA was found to be the most effective RI among those we tested (Figure 1F and 1G), while for the no-heating group, both PVSA and protein-based RIs showed similar results, effectively protecting RNA degradation (Figure S6A). PVSA also has no measurable negative effect on the downstream enzymes such as protease or reverse transcriptase (RTase), whereas the nucleobase-reactive chemical alternative, diethylpyrocarbonate (DEPC), does (Figure S6B)^22^. Each component was further confirmed by qPCR quantification using nuclei extracted from heavily fixed mouse brain tissue (Figure S7; STAR Methods).

### FX-seq for PFA-fixed and FFPE mouse brains

To achieve highly scalable sequencing of FX-treated nuclei, we employed combinatorial cell barcoding according to the updated sci-RNA-seq3 protocol, with some minor modifications (STAR Methods)^23^. To validate the reproducibility and performance of FX-seq, mice were perfused and brain tissue was extracted and fixed in PFA for 48 hours at 4°C in triplicate. Fixed brains were subjected to the FX procedure. Isolated nuclei were then combinatorially barcoded and sequenced (n = 3, Figure 1H and 1I). As a result, FX-seq successfully unlocks masked transcriptomic information in fixed biological samples by increasing the yield of cDNA synthesis, which was limited in PFA-fixed tissues. Specifically, FX-seq improved the data quality in terms of unique molecular identifiers (UMIs) and gene counts, revealing distinct cell types by recovering significantly more single-nucleus transcriptome information than untreated, Tris-buffered, and cat. 2-treated controls (n = 7,645, 8,815, 8,815, and 9,344 nuclei from three brains, respectively; Figure 1J–1N). Despite our optimized nucleus isolation procedure that preserves RNA quality, sequencing results from nuclei from PFA-fixed samples showed limited cell type identification due to poor RT efficiency (Figure S8A–S8H). However, FX-seq significantly improved cell type identification by enabling accurate classification of neurons, particularly distinguishing between medium spiny neurons (MSNs) and inhibitory interneurons. In both stand-alone and merged UMAP clustering, FX-seq facilitated a more accurate classification of specific cell types, including Purkinje cells, olfactory ensheathing cells (OEC), and macrophages, based on classical molecular classification criteria (Figure 1O–1T and S8I–1L; Table S1 and S2).

For comparison, we also performed the conventional snRNA-seq using freshly extracted nuclei with very short fixation (0.1% PFA for 10 min) employing a recent protocol from EasySci^24^ with minor modifications (STAR Methods). Although freshly extracted nuclei showed a higher transcript coverage, FX-seq of heavily fixed nuclei generated a dataset with comparable UMI and gene counts (Figure 2A and 2B). The lack of enzymatic digestion in fresh nuclei results in a higher cytoplasmic fraction and incomplete nucleus segregation (Figure S9A and S9B), which is corroborated by higher mitochondrial and ribosomal RNA reads as well as higher exonic read ratio (Figure 2C–2E). Nevertheless, the same cell types independently annotated in each dataset (Figure 2F and 2G; Table S3 and S4) were co-clustered in unsupervised clustering using the merged gene matrix (Figure 2H and 2I). The intra-cluster cellular distribution between fresh versus fixed and FX-treated is attributed to exonic read ratios (Figure 2J), which is further supported by a contrasting exonic read ratio of differentially expressed genes from each sample within identical clusters (Figure 2K). Overall, the FX treatment restores complex cellular resolution in heavily fixed tissues analogous to freshly extracted nuclei.

**Figure 2.**
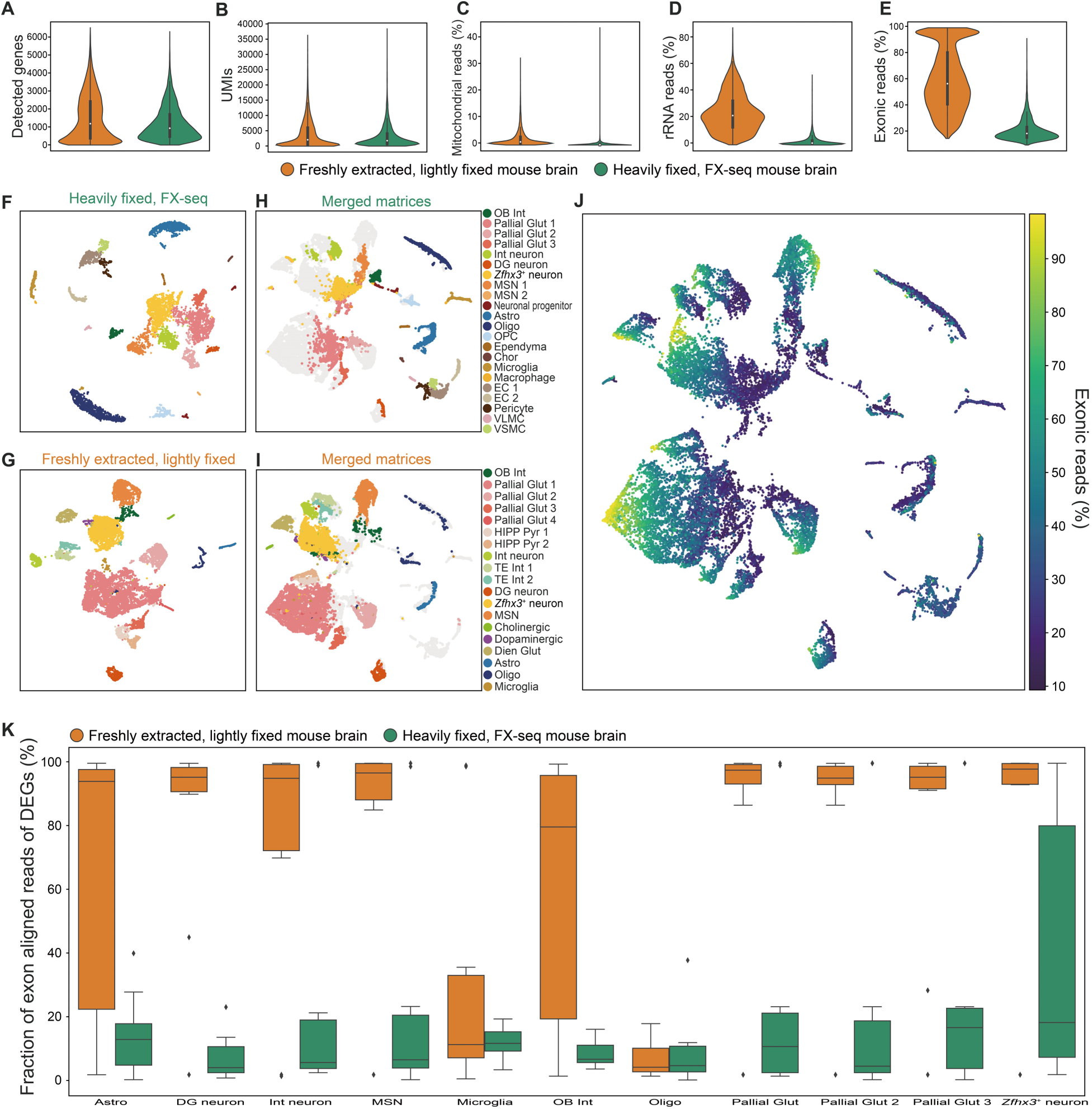
FX-seq of perfused, heavily fixed mouse brain tissue shows comparable performance to snRNA-seq of freshly extracted and lightly fixed mouse brain. (A–E) QC metrics of freshly extracted, lightly fixed mouse brain nuclei (n = 11,807) and FX-treated heavily fixed mouse brain (n = 7,569). The numbers of detected genes (A) and UMIs (B). Both were high in freshly extracted samples. The ratio of reads mapped to mitochondrial genes (C), rRNA (D), and exonic regions (E). High proportions of mitochondrial genes, rRNA, and exonic regions indirectly indicate contamination of cytoplasmic contents due to limited tissue digestion in fresh samples. (F–I) The UMAP analysis of each sample after QC cutoff (UMIs from 200 to 40,000 and mitochondrial reads < 1%) (F and G). Individual clusters were annotated based on the molecular criteria. In merged clusters, clusters with the same annotations in separate clustering are co-clustered, while the displayed distribution shows some shifts in the UMAP (H and I). (J) Color map of the percent of fractions mapped to exonic regions in the UMAP analysis of the merged gene expression matrix. The distribution shift between fresh and FX samples in the merged matrix is attributed to the degrees of their exonic reads fractions. (K) Distribution of the fraction of exonic reads in top 10 genes identified by differentially expressed gene analysis between fresh and FX-derived transcriptomes in the same cell types. Highly expressed genes in fresh samples have higher exonic read fractions, while highly expressed genes in FX samples have lower exonic read fractions. Abbreviations are as follows: OB Int, olfactory bulb interneuron; Pallial Glut, pallial glutamatergic neuron; Int neuron, interneuron; DG neuron, dentate gyrus neuron; MSN, medium spiny neuron; Astro, astrocyte; Oligo, oligodendrocyte; OPC, oligodendrocyte precursor; Ependyma, ependymal cell; Chor, choroid plexus epithelial cell; EC, endothelial cell; VLMC, vascular leptomeningeal cell; VSMC, vascular smooth muscle cell; HIPP Pyr, hippocampus pyramidal neuron; TE Int, telencephalon interneuron; Cholinergic, habenula cholinergic neuron; Dopaminergic, dopaminergic neuron; Dien Glut, diencephalic glutamatergic neuron.

We further investigated the application of FX-seq to mouse-derived FFPE samples with the goals of extending its utility and simulating clinical human FFPE samples. FX treatment was similarly performed on a triplicate of mouse brain FFPE tissue (Figure S10A; STAR Methods). Unfortunately, the lack of perfusion prior to embedding affected the sequencing results and compromised the overall quality of the obtained transcriptome data. Despite the reduced sample quality resulting from FFPE processing, FX-seq increased both the total number of UMIs and the number of genes detected (Figure S10B–S10D). Importantly, FX treatment provided additional information that improved cell type annotation in both stand-alone and merged UMAP clustering, clearly resolving specific cell populations such as pallial glutamatergic cells (expressing *Satb2*), *Zfhx3*^+^ neurons representing non-pallial neurons^25,26^, and inhibitory interneurons (Figure S10E–S10L and S11; Table S5 and S6). In contrast, the sequencing results from untreated control samples did not provide sufficient information to distinguish these specific cell types, resulting in their classification as generic neurons.

### Application to mouse brain and human clinical FFPE and H&E sections

In the next phase of our study, we investigated the compatibility of FX-seq with sectioned FFPE mouse brain and hematoxylin and eosin (H&E) stained sections. With minor protocol modifications, we successfully isolated up to 200,000 nuclei from a single 10 μm FFPE or H&E mouse brain section (Figure 3A and S5C and S5D; STAR Methods). However, sectioning of FFPE samples resulted in a reduction in total UMIs and gene counts, likely due to the physical cutting of nuclei. Despite this reduction, we observed similar clusters compared to the FFPE block experiments, indicating preserved transcriptomic information (Figure 3B and 3C). Some discrepancies between the transcriptomic signature between the FFPE block and sectioned samples may be attributed to the lack of mechanical homogenization of the sections, which ultimately leads to a shift in the cell population distribution. When FX-seq was applied to H&E-stained sections, mitochondrial and ribosomal RNA reads increased, while UMIs and gene counts decreased (Figure 3D and 3E). These observations may be due to limited enzymatic digestion efficiency and the effects of harsh staining conditions, leading to an accumulation of higher cellular debris (Figure S12A–S12D). Nevertheless, merged transcriptome analysis of FFPE and H&E sections revealed co-clustering of separately identified cell types, suggesting reliable and consistent transcriptomic profiles across both platforms (Figure 3F–3K and S12E–S12J; Table S7 and S8).

**Figure 3.**
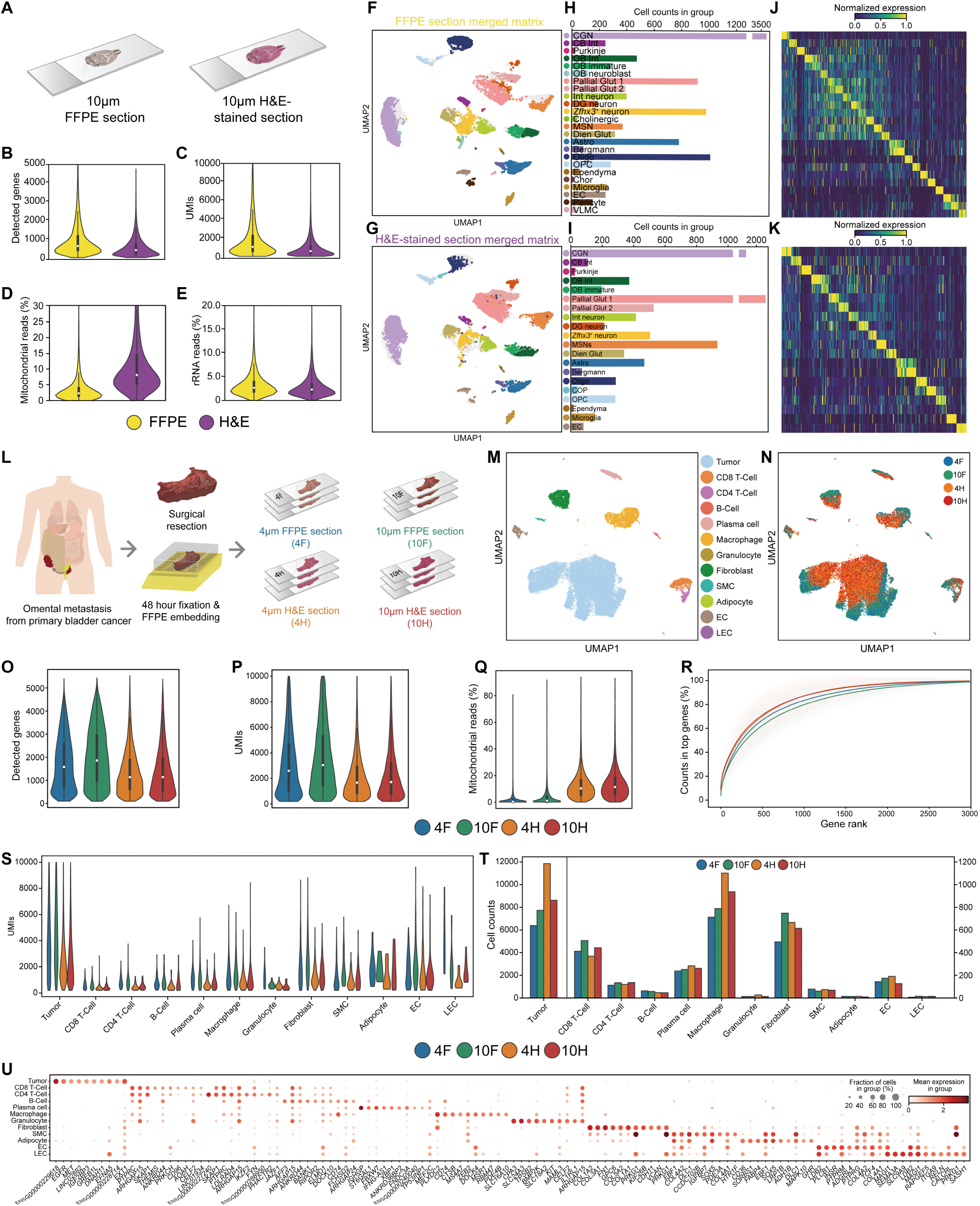
Application of FX-seq on thin FFPE sections with and without H&E staining. Compatibility testing of FX-seq with clinical settings. Nuclei were isolated from mouse brain sections and human omental metastasis cancer tissue with and without H&E staining. (A) Nuclei isolated from 10 μm FFPE and H&E sections of mouse brain FFPE blocks were analyzed by FX-seq. (B–E) QC metrics of the FFPE section (n = 11,334) and the H&E-stained section (n = 8,648) from (A). The number of detected genes (B), the number of UMIs per nucleus (C), the ratio of mapped reads to mitochondrial genes (D), and the ratio of mapped reads to rRNA (E). (F and G) The UMAP analysis of the FFPE section and the H&E-stained section. In the UMAP analysis of the merged gene expression matrix, the nuclei isolated from different conditions showed co-clustering of the same cell types. Although each cluster was annotated separately, the analysis revealed similarities in the gene expression patterns of the corresponding cell types across the different conditions. (H and I) Cell type annotation and cell number distribution of each cluster. (J and K) Specific expression of representative marker genes in each cluster. (L) Application of FX-seq to human nuclei from omental metastasis cancer samples prepared from 4 μm and 10 μm FFPE and H&E sections (labeled as 4F, 10F, 4H, and 10H, respectively) and analysis results are shown in (M to U, n = 8,590 from 4F, n = 10,395 from 10F, n = 14,658 from 4H, and n = 11,206 from 10H). (M) The UMAP analysis of the merged gene expression matrix of four conditions. (N) The distribution of individual nuclei in the UMAP (M), labeled according to the experimental conditions, shifts primarily according to H&E staining. (O–R) QC metrics of the FX-seq from (L). The number of detected genes (O), the number of UMIs per nucleus (P), the ratio of mapped reads to mitochondrial genes (Q), and the number of accounted UMIs along the top genes sorted by the number of detected UMIs in each condition (R). 4F had fewer genes and UMIs than 10F due to physical subsampling, with H&E staining having a greater effect than section thickness. (S) Distribution of UMIs in each cell type labeled by experimental condition. (T) Distribution of cell numbers for each cell type. (U) Dot plot of marker genes specific to each cluster. Abbreviations are as follows: CGN, cerebellar granule neuron; CB Int, cerebellum interneuron; Purkinje, Purkinje cells; OB Int, olfactory bulb interneuron; OB immature, olfactory bulb immature neuron; OB neuroblast, olfactory bulb neuroblast; Pallial Glut, pallial glutamatergic neuron; Int neuron, interneuron; DG neuron, dentate gyrus neuron; Cholinergic, habenula cholinergic neuron; MSN, medium spiny neuron; Dien Glut, diencephalic glutamatergic neuron; Astro, astrocyte; Bergmann, Bergmann glia; Oligo, oligodendrocyte; Ependyma, ependymal cells; Chor, choroid plexus epithelial cell; EC, endothelial cell; VLMC, vascular leptomeningeal cell; COP, committed oligodendrocyte precursor; OPC, oligodendrocyte precursor; SMC, smooth muscle cell; LEC, lymphatic endothelial cell.

We further investigated whether the section thickness and H&E staining affect the quality of sequencing data from human clinical samples. FX-seq was applied to a surgical specimen excised from a metastatic omental mass originating from primary bladder cancer (surgical specimen dimensions: approximately 2.5 cm by 1.2 cm per slide) (Fig. S13A and S13B; STAR Methods). The specimen was immediately fixed with 4% PFA for 48 hours at 4°C and subjected to FFPE processing. The FFPE block was sectioned and H&E stained at two different thicknesses, 4 μm and 10 μm, which are typically employed for pathologic examination (Figure 3L). The resulting sections were analyzed using the FX-seq protocol modified for FFPE sections. As expected, the transcriptomes of single nuclei obtained from both FFPE and H&E sections were consistently co-identified based on their cell types (Figure 3M and 3N), regardless of decreased UMIs and increased mitochondrial counts due to staining and thin sectioning (Figure 3O–3R). It should also be noted that the recovered number of UMIs and cells per population in each experimental group had similar characteristics as a consolidated group (Figure 3S–3T). Collectively, these results suggest that FX-seq, as a scalable snRNA-seq method, enabled the recovery of a high-quality single-nucleus–level transcriptome from FFPE and H&E sections of human surgical specimens (Figure 3U; Table S9).

### Scalability of FX-seq unravels unique molecular pathways associated with TKI resistance

Large-scale transcriptome profiling at single-cell resolution provides better coverage of cell types, leading to the identification of diverse and rare cell types for a better understanding of biological processes and cellular heterogeneity^23,27,28^. To assess the scalability and versatility of FX-seq, we studied an FFPE block of a gastrointestinal stromal tumor (GIST) undergoing palliative surgery due to recent tumor progression after a partial response to 5 years of tyrosine kinase inhibitor (imatinib) treatment. The tumor originated from the small intestine and measured approximately 1.5 cm x 0.8 cm x 1 cm in size (Figure 4A and S14A and S14B). In a single experiment, we isolated approximately 4 million nuclei from one-tenth of a homogenized GIST FFPE block and profiled a subset of 199,293 nuclei with an average of approximately 4,600 reads per nucleus (STAR Methods). The resulting raw sequencing data was further processed for quality control and doublet removal, resulting in a total of 171,992 single-nucleus transcriptomes with an average UMI count of 1543.7 (Figure 4B–4F; STAR Methods). The analyzed dataset shows a comprehensive classification of diverse cell types, including rare immune cell populations such as conventional dendritic cells (cDC) and plasmacytoid dendritic cells (pDC) (Figure 4G; Table S10 and S11).

**Figure 4.**
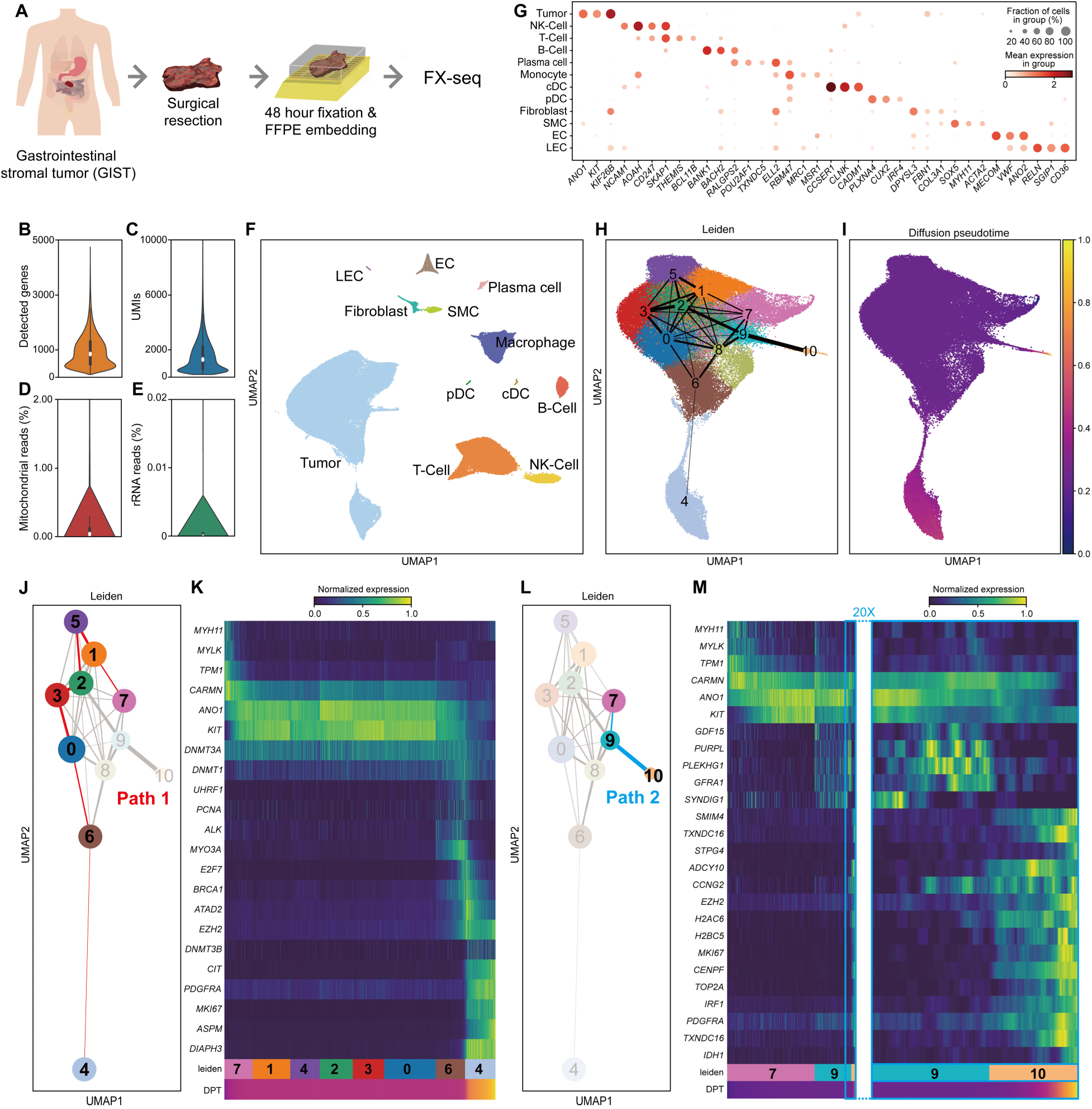
Scalable FX-seq analysis of human surgical specimen from gastrointestinal stromal tumor (GIST) FFPE block. (A) Surgically resected human GIST tissue was immediately immersed in 4% PFA, postfixed for 48 hours at 4°C, and embedded in an FFPE block. Nuclei were isolated from the FFPE block and labeled combinatorially using sci-RNA-seq3 for snRNA-seq library preparation. Transcriptomes from 171,992 nuclei were recovered after QC and doublet removal. (B) Dot plot of marker genes specific to each cluster. (C–F) QC metrics of the FX-seq from (A). The number of detected genes (C), the number of UMIs per nucleus (D), the ratio of mapped reads to mitochondrial genes (E), and the ratio of mapped reads to rRNA (F). Nuclei isolated from the FFPE block showed minimal contamination with mitochondrial genes and rRNA. (G) The UMAP analysis of transcriptome distribution and clustering. Various cell types were identified based on molecular criteria. (H and I) Constructed cluster networks based on PAGA analysis (H) and their diffusion pseudotime heatmap (I). (J–M) Resolved transcriptome paths and their gene expression patterns. Resolved tumor path 1 (J) and its corresponding gene expression changes per cluster (K). Resolved tumor path 2 (L) and its corresponding gene expression changes per cluster (M), the right blue frame image is a 20x magnified expression profile change in path 2, from tumor subcluster 9 to 11. Abbreviations are as follows: LEC, lymphatic endothelial cell; EC, endothelial cell; SMC, smooth muscle cell; pDC, plasmacytoid dendritic cell; cDC, classical dendritic cell; NK cell, natural killer cell; DPT, diffusion pseudotime.

Among the cells in the dominant cluster representing tumor populations, we found a significant increase in the expression of *ANO1*, a specific marker gene associated with GIST tumor cells and interstitial cells of Cajal (ICC), recognized as GIST progenitor cells^29,30^ (Figure S15A). Interestingly, we identified separate clusters characterized by significant expression of the driving oncogenes KIT and PDGFRA (Figure S15A), suggesting that the expression levels of these genes are independently altered during the development of GIST from ICC^31,32^. Moreover, only the PDGFRA^high^ population expressed cell proliferation markers such as *CENPF*, *TOP2A*, and *MKI67* (Figure S15B). Based on the reconstructed trajectories using pseudo-time inference and partition-based graph abstraction (PAGA) analysis^33^ of the tumor cluster (Figure 4H and 4I), we found two distinct paths of transcriptional shift showing a transition from the normal ICC [expressing *MYH11*, *MYLK*, and *TPM1*^34^ (Figure S15C)] to the proliferating PDGFRA^high^ cells via non-proliferative KIT^high^ cells.

We hypothesize that these two pathways leading to proliferating populations may contain transcriptomic information relevant to imatinib-resistant cells in GIST with a *KIT* mutation (p.N822K), which is commonly observed as a secondary mutation associated with acquired resistance developed after long-term imatinib treatment^35,36^. Along path 1 (Figure 4J), the transcriptomes show transiently increased expression of DNA methylation-related genes, including *PCNA*, *DNMT1*, *UHRF1,* and sustained upregulation of the histone methylation enzyme encoded by *EZH2.* Additionally, transient expression of several genes, including *ALK, ATAD2, and MYO3A,* was observed (Figure 4K and S16). We further hypothesize that these transient gene expression events, along with epigenetic modifications, may contribute to imatinib resistance and result in altered transcriptome patterns of proliferating *PDGFRA*^high^ cells represented by overexpression of *DNMT3B, CIT,* and *DIAPH3*. Along path 2 (Figure 4L), we observed a similar overexpression of EZH2 without changes in DNA methylation-related genes. Notably, in the non-proliferative tumor population (Cluster 9 in Figure 3L), we observed transiently increased gene expression of *GDF15*, *PURPL*, *PLEKHG1*, *GFRA1*, and *SYNDIG1*, which was subsequently decreased at the end of path 2. In the proliferative population, we observed abnormal gene expression, including the expression of germline-specific genes such as *SPAG1*, *SPTG4*, *ADCY10*, and *CCNG2*, along with proliferation markers (Figure 4M and S17).

In summary, our observations suggest that the two distinct paths, characterized by abnormal gene expression patterns and transient gene expression changes, provide valuable insights into the cellular response and acquisition of resistance to imatinib. Path 1 involves gene expression changes related to DNA methylation, which is consistent with previous reports of aberrant DNA methylation in imatinib resistance in chronic myelogenous leukemia (CML)^37,38^. Path 2 shows the expression of germline-specific genes that are typically not expressed in other tumor and normal cells, representing a putative rare cell population with a de novo path to imatinib resistance identified by large-scale snRNA-seq. Overall, we have demonstrated the high scalability and data quality of FX-seq analysis of clinical FFPE samples, and showcased a new opportunity to discover novel molecular markers to help understand the dynamic nature of integrated cellular processes associated with human disease.

### Integration of pathological diagnosis and snRNA-seq

Lastly, we employed FX-seq to explore the single-nucleus transcriptome of archived human colorectal cancer (CRC) FFPE slides stored for approximately 2 months. Based on pathologist annotation, identified tumor (T) regions were physically separated from non-tumor (NT) regions of FFPE sections. (Figure S18A–S18J). For each sample, we performed FX treatment, isolated nuclei, and barcoded them using 1st RT primers based on their regional origin, and the resulting nuclei were subjected to downstream combinatorial labeling and sequencing library construction (Figure 5A; STAR Methods). This approach allows us to perform FX-seq with spatially contextualized pathological annotation.

**Figure 5.**
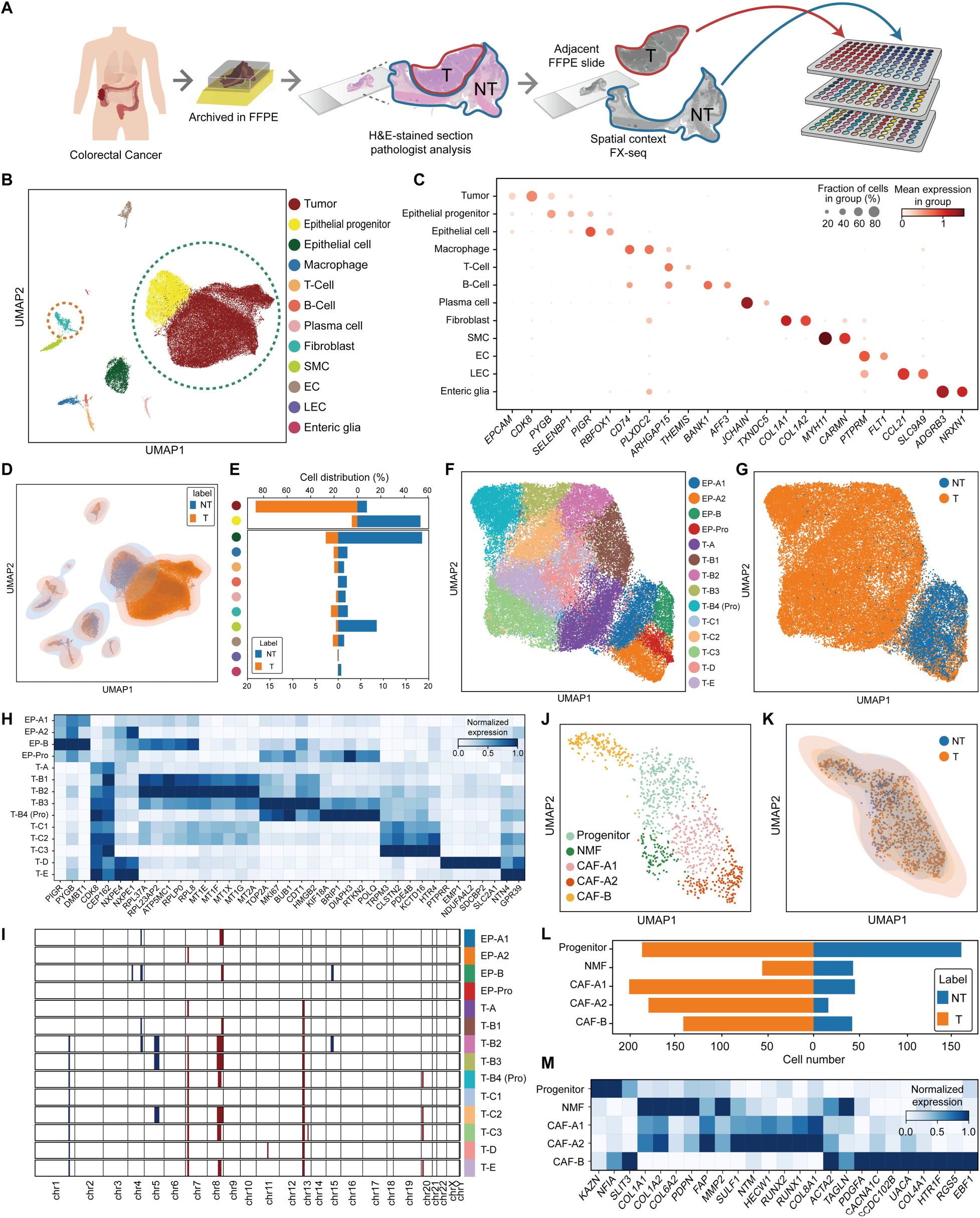
FX-seq analysis of the archived human colorectal cancer (CRC) FFPE specimen with spatial annotations. (A) Schematic procedure of FX-seq on the archived CRC FFPE specimen. The FFPE section was annotated by a pathologist to distinguish tumor (T) and non-tumor (NT) regions in H&E sections. Adjacent 10 μm FFPE sections were scraped, and nuclei were isolated for FX-seq with cell labeling by the first step of 3-stage combinatorial labeling (sci-RNA-seq3) to differentiate their regional origin (T vs. NT). (B) The UMAP analysis of the transcriptomes labeled by regional origin from FFPE sections (n = 47,966 from T and n = 13,559 from NT). (C) Representative marker genes for each cluster. (D) Distribution of cell populations labeled by regional origin in (B). (E) The ratio of cell numbers belonging to each cluster was calculated from all cells identified in each region. Colored dots represent the same cell type annotation as shown in (B). (F) The UMAP analysis to identify the subcluster of the merged gene matrix of epithelial progenitor and tumor populations shown in the green dotted circle in (B). (G) Distribution of cell populations labeled by regional origin in (F). The progenitor population was identified as having a mixed distribution of nuclei from the T and NT regions, whereas the tumor population was specific for nuclei from the T region. (H) Representative marker genes of cell types identified by sub-cluster analysis in (F). (I) Cluster-level CNV inference results from the gene expression of the tumor and epithelial progenitor populations using the inferCNV package. (J) The UMAP analysis to identify the subcluster of the fibroblast cluster shown in the orange dotted circle in (B). (K) Population distribution of the fibroblast subclusters labeled by regional origin in (J). (L) Cell number distribution of each cluster shown in (J and K). (M) Representative marker genes identified by subcluster analysis in (J). Abbreviations are as follows: SMC, smooth muscle cell; EC, endothelial cell; LEC, lymphatic endothelial cell; Enteric glia, enteric glial cell; EP, epithelial cell; Pro, progenitor cell; NMF, normal fibroblast; CAF, cancer-associated fibroblast;

The sequencing results reveal the presence of canonical cell types residing in CRC, including immune cells, endothelial cells, enteric glia, and connective cells with a predominant population of epithelial lineage cells (Figure 5B and 5C; Table S12). Distilled regional barcodes encoding the spatial origin of individual nuclei allow identification of their regional distribution within each cluster (Figure 5D and 5E). Interestingly, the large epithelial lineage clusters, which are distinct from normal epithelial cells, are identified as epithelial progenitor cells and tumor cells based on their regional origin (Figure 5F and 5G). We cross-validated this assignment using the molecular criteria of loss of polymeric immunoglobulin receptor (*PIGR*) expression^39^ and gain of *CDK8*^40^ (Figure 5C and S19). Subcluster analysis of the cluster in Figure 5G allows the elucidation of distinct transcriptome profiles in each population assigned to progenitor and tumor cells (Figure 5H; Table S13). Application of the inferCNV package^41^ to our dataset provides insights into changes in the copy number variation (CNV) states in both epithelial progenitor and tumor cells compared to the normal epithelial cell population. Similar to previously reported scRNA-seq of CRC samples^42^, a significant gain of Chr7p, Chr8q, Chr13q, and Chr 20p and loss of Chr1.p are observed in tumors, whereas the epithelial progenitor cells show minimal change in the CNV state (Figure 5I and S20). The CNVs observed in our study correspond to a subset of previously reported CNVs derived by scRNA-seq. This finding confirms that FX-seq can provide high-quality data from clinical specimens.

FX-seq also provides insight into the complex tumor microenvironment (TME) in CRC tissue. Additional subclustering analysis of the fibroblast population reveals two distinct cancer-associated fibroblasts (CAFs), an activated subset of fibroblasts that act as key regulators in the TME (Figure 5J; Table S13). Incorporating the regional annotation into the fibroblast population consolidates our cell type annotation, as CAF populations are biased toward tumor regions, whereas normal fibroblasts (NMF) and their progenitors show an even distribution in both tumor and non-tumor regions (Figure 5K and 5L). Two distinct CAF populations (CAF-A and CAF-B) are identified based on their transcriptomic states, and specific genes enriched in each population were identified (Figure 5M). The analysis of cluster-specific genes, including *FAP*, *PDPN*, *COL1A2*, and *MMP2* for CAF-A, and *ACTA2*, *TAGLN*, and *PDGFRA* for CAF-B, shows a strong correlation with previous studies^43^. In addition, our analysis reveals additional marker genes that are more specific to each cluster, providing further insight into the heterogeneity of CAF populations. Altogether, FX-seq can link surgical pathology diagnoses to their associated transcriptomes, and further investigate cellular heterogeneity in the TME.

Based on our observations, we further classified the identified tumor population as intrinsic consensus molecular subtype 2 (iCMS2), which is a subtype for malignant epithelial cells based on the scRNA-seq analysis of CRC and is characterized by notable copy number alterations^42^. Furthermore, the increased expression of *LGR5* in the tumor population further supports our classification of iCMS2, which is likely to arise by expansion from *LGR5*^+^ crypt-bottom stem cells^42^ (Figure S19). iCMS2 is reported to be predominantly microsatellite stable (MSS) and does not harbor mutations in *KRAS*, which is consistent with the diagnostic result of MSS– and *KRAS*–wildtype in our case. Additionally, the detection of CAF populations in the TME indicates a fibrotic environment in the tumor. Taken together, we classified the tumor in our study as iCMS2_MSS_F according to the proposed IMF classification using intrinsic epithelial subtype (I), microsatellite instability status (M), and fibrosis (F)^42^. Our analysis demonstrates that FX-seq has the potential to serve as a diagnostic tool by obtaining single-nucleus transcriptomes from FFPE sections. This capability enables the identification of cancer subtypes that require information at the single-cell or single-nucleus level resolution, eliminating the need for additional surgical procedures to obtain fresh specimens. By employing FFPE sections generated from routine histology, FX-seq offers valuable new opportunities for clinical diagnosis.

## DISCUSSION

In this work, we developed FX-seq to enable single-nucleus level analysis of various heavily PFA-fixed and clinical FFPE samples by recovering transcriptome information. FX-seq demonstrated solid applicability in various biological and clinical contexts so we expect the widespread use of FX-seq to recover transcriptome with 1) snRNA-seq from PFA- or FFPE-preserved tissues for easy storage and transport, 2) snRNA-seq atlas for archived human clinical FFPE samples with clinical annotations. The study of human diseases and their pathological markers with long-banked FFPE clinical tissues will provide new diagnostic criteria for precision medicine and will open new opportunities for drug treatment planning by examining features found in various disease samples through incorporating analytical and experimental tools for studying druggable targets^44^.

Animal studies for snRNA-seq analysis could also be greatly improved by simply perfusing the animals with PFA to permanently protect and fix the transcriptome. This feature eliminates the need for extensive efforts for sampling from fresh or fresh-frozen tissues and will significantly improve the reproducibility of the measurement by reducing unwanted RNA degradation and stress-induced gene expression during tissue digestion of fresh tissues to prepare single-cell suspensions^45^. Currently, cells and tissues with significant amounts of RNases in subcellular organelles cannot be analyzed by the conventional droplet-based labeling approach due to the severe degradation of RNAs after cell lysis in the droplet. However, these cells with endogenous RNases can be analyzed by the FX-seq protocol using PFA fixation and PVSA, so that the endogenous RNases are effectively inactivated by the FX-seq process. Finally, cells of pathogenic origin can also be analyzed by FX-seq due to the safe inactivation process by PFA fixation^46^.

### Limitations of the study

Despite the potential of FX-seq for high-resolution analysis of clinical FFPE samples, its current application is limited to specific conditions and has not been extensively explored. First, our demonstration of FX-seq was primarily focused on samples with relatively higher RNA quality, as library construction yields may be insufficient for highly degraded FFPE samples. Second, FX-seq has only been tested in combinatorial barcoding schemes and requires further validation for compatibility with other commercial platforms such as 10X chromium. Regardless of these limitations, we envision that FX-seq will facilitate highly scalable, effective single-nucleus transcriptome analysis from human clinical FFPE samples and various animal models using PFA fixation.

## ACKNOWLEDGEMENTS

We thank past members of the Sohn laboratory who contributed to protocol development (Hyunjun Ju, Junhee Kim, and Soyeon Kim). We also thank Taeyun Ku, In-Kyung Jung, Jihwan Park, and Jong-Eun Park who provided critical comments on our manuscript. **Funding:** C.H.S. was supported by grants from Institute for Basic Science (IBS-D01-026), National Research Foundation (NRF) of Korea, funded by the Ministry of Science and ICT (2021R1C1C1004092 and 2021H1D3A2A02096517), Yonsei University (2022-22-0093), Korea Foundation for Women in Science, Engineering, and Technology (WISET) (2023-277), and Illumina Asia-Pacific and Japan.

## AUTHOR CONTRIBUTIONS

C.H.S. and H.-E.P. conceived FX-seq. H.-E.P. and Y.T.L. developed techniques with assistance from H.J., J.L., and J.W.L.; H.-E.P. and Y.T.L. performed experiments with assistance from J.W.L.; J.L. and C.W.L. synthesized crosslinkers.; E.K. synthesized catalysts during the early phase of the study.; Y.-L.S. prepared purified Tn5 transposase with assistance from J.K.H.; H.-E.P. performed computational analyses with assistance from Y.T.L., S.-Y.K., Y.D.H., and H.K.; Y.D.H., and H.K. provided clinical specimens with pathological annotations.; H.-E.P, Y.T.L., and C.H.S. wrote the manuscript with input from all coauthors; and C.H.S. supervised the project.

## DECLARATION OF INTERESTS

Patent applications have been filed by Yonsei University related to this work.

## INCLUSION AND DIVERSITY

We support inclusive, diverse, and equitable conduct of research.

## STAR★METHODS

### KEY RESOURCES TABLE

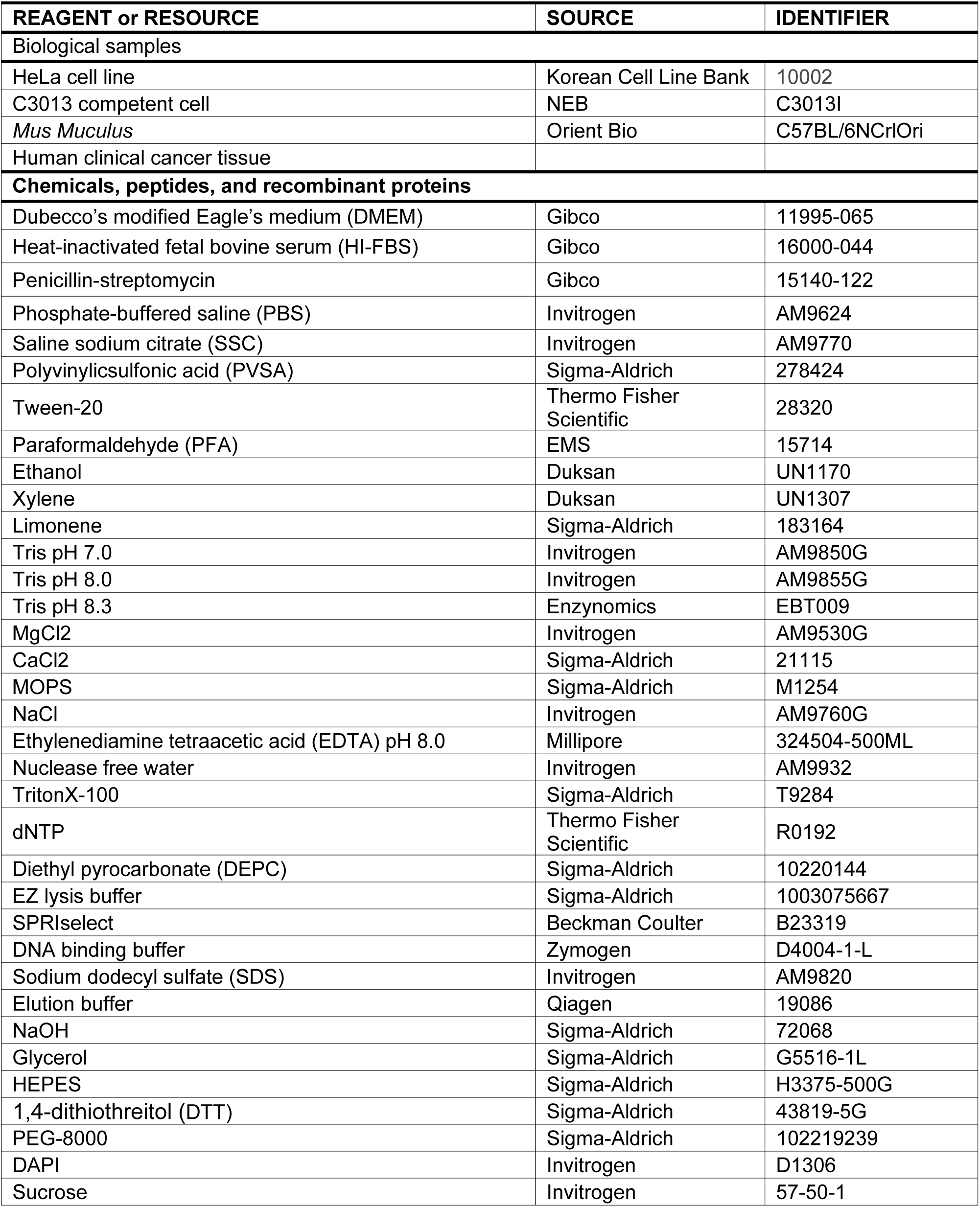
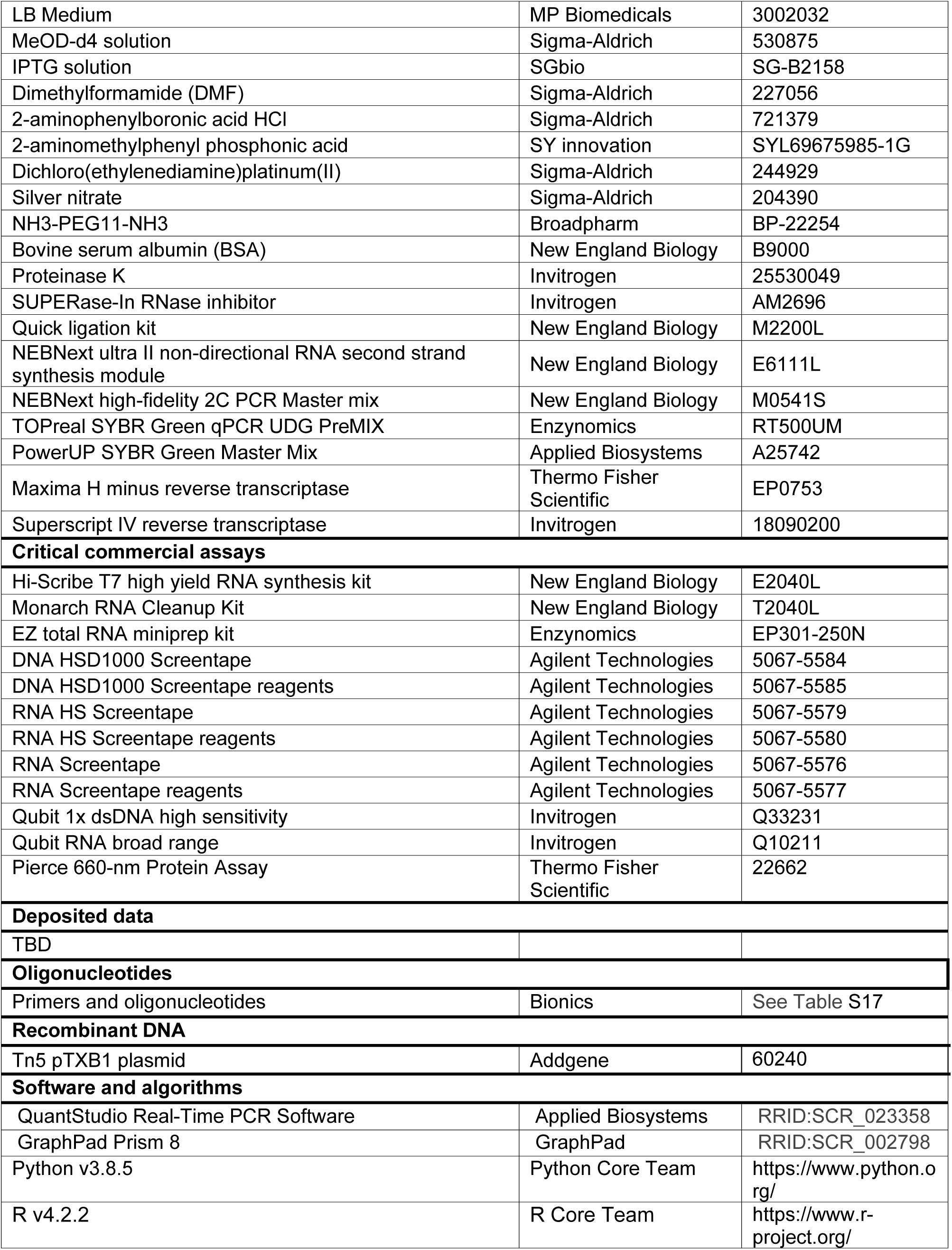
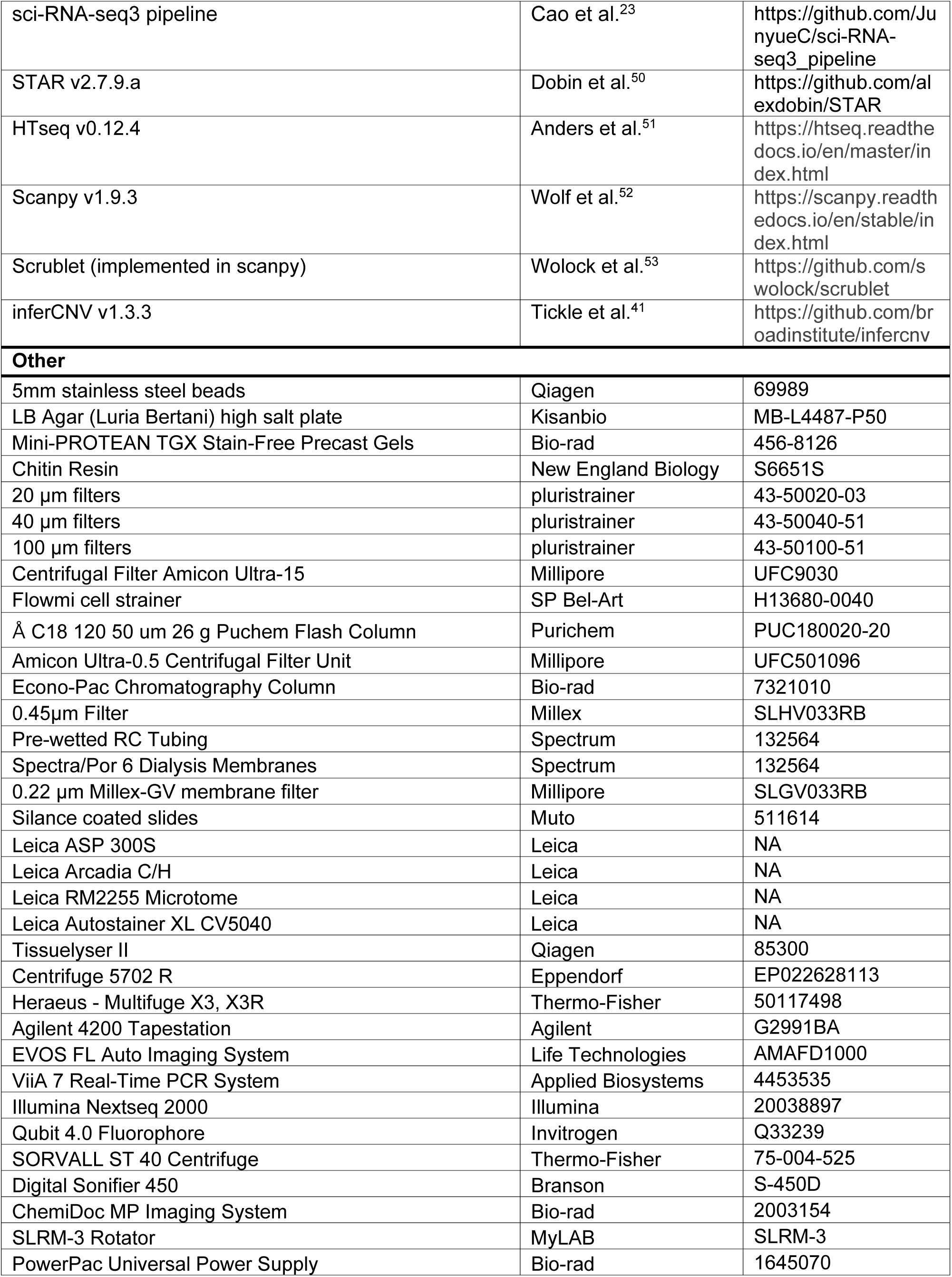
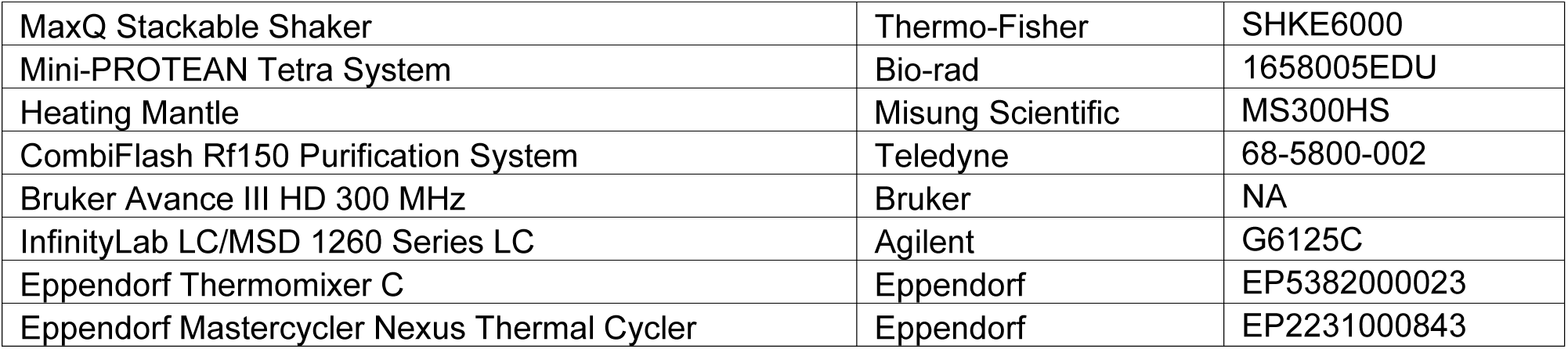

### RESOURCE AVAILABILITY

#### Lead contact

Further information and requests for resources and reagents should be directed to and will be fulfilled by the lead contact, Chang Ho Sohn.

#### Materials availability

All unique/stable reagents generated in this study are available from the lead contact with a completed Materials Transfer Agreement.

#### Data and code availability

The data generated in this study has been submitted to NCBI Gene Expression Omnibus in both raw and processed forms. The accession number will be shared as soon as it becomes available.

This paper does not report the original code. Any additional information required to reanalyze the data reported in this paper is available from the lead contact upon request.

### EXPERIMENTAL MODEL AND SUBJECT DETAILS

#### HeLa cell culture

HeLa cells were cultured in Dubecco’s modified Eagle’s medium (DMEM) (Gibco) supplemented with 10% heat-inactivated fetal bovine serum (HI-FBS) (Gibco) and 1% penicillin-streptomycin (Gibco). Cells were incubated at 37°C in 5% CO_2_ and split three times per week.

#### Animals

All animal handling and experiments were performed under the guidelines of the Institutional Animal Care and Use Committee of Yonsei University. 8-week-old C57BL/6N wild-type mice were purchased from Orient Bio, South Korea, and maintained under strict SPF conditions at the animal facility of the Yonsei Advanced Science Institute. Mice were sacrificed between 8–10 weeks of age.

#### Human subjects

All human patient samples utilized in this work were sanctioned through preoperative written consent within the borders outlined by Yonsei University Severance Hospital Institutional Review Board.

### METHOD DETAILS

#### Confirmation of PFA inhibition of *in vitro* isolated RNA

All *in vitro* transcription of isolated RNA was performed by Hi-Scribe T7 high yield RNA synthesis kit (NEB) following manufacturer’s protocols. *In vitro* transcribed RNA was purified by adding 1 volume of SPRIselect (Beckman Coulter) and eluting by following the manufacturer’s protocols. The purified isolate RNA was then fixed (50 ng/μL purified single *PHF11* and *SET7/9* RNA from *in vitro* transcription, 4% PFA in 1X PBS) by shaking at 37°C for varying time intervals. Fixed RNA was purified using column purification following the manufacturer’s protocols and quantified using a Qubit RNA BR kit (Invitrogen). Effects of PFA inhibition during reverse transcription was assessed by reverse transcription and qPCR quantification. Reverse transcribing of the fixed RNA was performed as follows: 2 μL of 0.1 ng/μL fixed RNA, 2 μL of 5X Maxima H Minus Reaction buffer, 0.25 μL Maxima H Minus reverse transcriptase, 0.25 μL SUPERase-In, 1 μL of 10 μM reverse primer, 0.5 μL of 10mM dNTP, 2 μL nuclease-free water. The reverse transcribed cDNA level was quantified following the qPCR protocol described below.

#### *in vitro* isolated RNA PFA crosslinking removal

*In vitro* transcribed RNA was fixed (200 ng/μL purified single *PHF11* RNA from *in vitro* transcription, 4% PFA in 1X PBS) at room temperature for 16 hours. Fixed RNA was purified using column purification following the manufacturer’s protocols and quantified using a Qubit RNA BR kit (Invitrogen). The degree of crosslinking removal was tested by treating the fixed RNA in organocatalyst solution (10 ng/μL fixed RNA, 10 mM Tris pH 7.0, 10 mM cat. 1 or cat. 2, 0.1% v/v SUPERase-In) for 30 minutes at 55°C. Treated RNA was purified by adding 1 volume of SPRIselect (Beckman Coulter) and eluted following the manufacturer’s protocols. The purified RNA was then quantified and reverse transcribed as follows: 2 μL of 0.1 ng/μL *PHF11* RNA, 2 μL of 5X Maxima H Minus Reaction buffer, 0.25 μL Maxima H Minus reverse transcriptase, 0.25 μL SUPERase-In, 1 μL of 10 μM reverse primer, 0.5 μL of 10 mM dNTP, 2 μL nuclease-free water. The reverse transcribed cDNA level was quantified following the qPCR protocol described below.

#### HeLa total RNA PFA crosslinking removal

Total RNA was extracted from fresh cultured HeLa cells using column purification methods following the manufacturer’s protocol. Extracted fresh total RNA was fixed (200 ng/μL purified total RNA, 4% PFA in 1X PBS) at room temperature for 16 hours. Fixed RNA was purified using column purification following the manufacturer’s protocols and quantified using a Qubit RNA BR kit (Invitrogen). The degree of crosslinking removal was tested by treating the fixed RNA in organocatalyst solution (10 ng/μL fixed RNA, 10 mM Tris pH 7.0, 10 mM cat. 1 or cat. 2, 0.1% v/v SUPERase-In) for 30 minutes at 55°C. Treated RNA was purified by adding 1 volume of SPRIselect (Beckman Coulter) and eluted following the manufacturer’s protocols. The purified RNA was then quantified and reverse transcribed as follows: 2 μL of 0.1 ng/μL total RNA, 2 μL of 5X Maxima H Minus Reaction buffer, 0.25 μL Maxima H Minus reverse transcriptase, 0.25 μL SUPERase-In, 1 μL of 10 μM oligo dT primer, 0.5 μL of 10 mM dNTP, 2 μL nuclease-free water. The reverse transcribed cDNA level was quantified following the qPCR protocol described below.

#### *in vitro* isolated RNA additional crosslinking

*In vitro* transcribed RNA was shaken at 37°C for 15 minutes [50 ng/μL purified *PHF11* RNA from *in vitro* transcription, 1X MOPS (1X PBS for PFA crosslinking) with 1–8% PFA or 0.5X–4X Pt crosslinker] (1X crosslinker is equivalent to 58.7 μM in all experiments for this work). Each sample was immediately placed on ice, diluted, and visualized on the Agilent 4200 Tapestation (Agilent) to assess the crosslinking degree. The reverse transcription solution was prepared as follows: 2 μL of 0.1 ng/μL crosslinked *PHF11* RNA, 4 μL of 5X SuperScript buffer, 0.5 μL SuperScript Reverse Transcriptase, 0.5 μL SUPERase-In, 1 μL of 0.1 M 1,4-dithiothreitol (DTT), 2 μL of 10 μM reverse primer, 1 μL of 10 mM dNTP, 9 μL nuclease-free water. After incubation at 55°C for 30 minutes, the degree of inhibition of each crosslinker was quantified using qPCR following the protocol described below.

#### HeLa cell line experiment

For HeLa cell line experiments, 1.4–2 million cells were collected and spun down at 200 g for 2 minutes at room temperature. Cells were washed once with 1X PBS (Gibco) and fixed in 4% PFA in 1X PBS solution for 15 minutes at 4°C. Cells were spun down at 200 g for 3 minutes at 4°C and washed with the wash buffer (1X PBS, 0.025% Tween-20). The washed cells were permeabilized in PBST solution (1X PBS, 0.05% TritonX-100, and 0.4 mg/mL BSA) for 15 minutes at room temperature. Cells were then washed in the wash buffer. Fixed cells were filtered using a 40 μm cell strainer (pluriStrainer) with a wide-bore tip on ice. Cells were counted and diluted to a final concentration of 5k cells/μL. The filtered cells were incubated in CL(–) solution (1X MOPS, 0.025% Tween-20) for 5 minutes in a cold room and repeated. Then CL(+) solution (1X MOPS, 0.025% Tween-20, 0.2X crosslinker) was added and shaken at 25°C for 15 minutes. Crosslinker concentrations were optimized for each sample type because the reaction kinetics varies significantly between ECM-free cell lines and ECM-rich tissues. Crosslinked cells were centrifuged at 200 g and resuspended in the wash buffer and counted. Cells were aliquoted to a final concentration of 500 cells/μL per experimental condition. Counted and aliquoted cells were washed with CT(–) solution (200 mM Tris pH 8.3). The CT(+) solution (200 mM Tris pH 8.3, 50 mM cat. 2) was applied and shaken at 55°C for varying time intervals. Samples were immediately placed on ice and centrifuged at 200 g. A portion of the supernatant was taken to measure the RNA concentration using the Qubit RNA BR assay (Invitrogen) according to the manufacturer’s manual. The rest of the supernatant was discarded, and the cell pellets were resuspended in PBSRI (1X PBS, 0.4 mg/mL BSA, 0.1% v/v SUPERase-In) and counted. The cells were reverse transcribed at 55°C for 30 minutes as follows: 11 μL 4,300 cell/μL, 4 μL of 5X SuperScript buffer, 1 μL SuperScript reverse transcriptase, 1 μL SUPERase-In, 1 μL of 10 μM reverse primer, 1 μL of 10mM dNTP, 1 μL DTT. The reverse transcribed cDNA level was quantified following the qPCR protocol described below.

#### qPCR screening

In this work, we optimized experimental conditions and sample conditions through qPCR screenings. Cell lines, whole tissues or tissue sections were crosslinked, reversed, nuclei isolated, and washed in a parallel procedure as described above. For *in situ* experiments, nuclei were directly lysed by adding an equal volume of 2X lysis mixture (20 mM Tris pH 8.0, 400 mM NaCl, 100 mM EDTA pH 8.0, 4.4% SDS, 3.34 mg/mL Proteinase K) to the reverse transcribed nuclei and gently suspending. The lysis reaction was performed using the following cycle setting: 55°C for 30 minutes, 85°C for 15 minutes. Reverse transcribed cDNA was purified by adding 1 volume of SPRIselect to the lysate and eluted following the manufacturer’s protocol. For *in vitro* experiments, the lysis step was omitted. For each well of the qPCR plate, 10 μL TOPreal SYBR Green qPCR UDG PreMIX (Enzynomics) or PowerUP SYBR Green Master Mix (Applied Biosystems), 1 μL of 10 μM reverse primer, 1 μL of 10 μM forward primer, 2 μL eluted cDNA, 6 μL nuclease-free water were mixed and 18 μL was aliquoted. Each tested condition was technically triplicated within each plate to maintain technical integrity. The qPCR reaction was performed by the ViiA 7 Real-Time PCR System (Applied Biosystems) using the following cycle setting: initial denaturation step at 95°C for 15 minutes; amplification step for 40 cycles at 95°C for 10 seconds, 60°C for 15 seconds, 72°C for 30 seconds (95°C for 15 seconds, 60°C for 1 minute for PowerUP SYBR Green Master Mix); final extension step at 72°C for 10 minutes; Melting curve measurement step at 95°C for 15 seconds, 60°C for 1 minute, 95°C for 30 seconds.

#### PFA fixation of mouse brain

To obtain PFA-fixed mouse brain tissue, 8–10 week-old mice were anesthetized (0.8 L/min oxygen with 3% isoflurane) and maintained (0.8 L/min oxygen with 2% isoflurane). Mice were then transcardially perfused with ice-cold 1X PBS (Gibco) followed by an ice-cold 4% PFA (EMS), 1X PBS (Gibco) solution. Perfused mice were decapitated to collect the whole brain. The dissected brain was placed in 4% PFA (EMS) in 1X PBS solution, and shaken in a cold room for 48 hours.

#### FFPE embedding of mouse brain

To obtain FFPE-embedded mouse brains, 8–10 week old mice were euthanized in a CO_2_ chamber. Mice were immediately decapitated, and their brains were dissected. The obtained tissue was placed in 4% PFA (EMS) in 1X PBS solution in a cold room, and shaken for 48 hours. The fixative solution was changed once after 2–3 hours of shaking to ensure consistent brain quality. After 48 hours, heavily fixed brains were placed on embedding cassettes and embedded in FFPE blocks according to the following protocol: dehydration with one change of 70% ethanol for 1 hour, one change of 80% ethanol for 1 hour, two changes of 95% ethanol for 1 hour each, three changes of 100% ethanol for 1 hour each, three changes of xylene for 1 hour each, three changes of heated (56–58°C) paraffin wax for 1 hour each, and finally embedded in paraffin blocks. Leica ASP 300S (Leica) equipment was used for tissue processing and Leica Arcadia C/H (Leica) equipment was used for paraffin embedding. All FFPE embedding protocols were performed at the Department of Histopathology, Avison Biomedical Research Center, Yonsei University.

#### FFPE embedding of human surgical samples

Human tumor samples were obtained from Yonsei Severance Hospital. All freshly obtained samples were immediately placed in 4% PFA (EMS) in 1X PBS solution, and shaken in a cold room for 48 hours. The fixative solution was changed once after 2-3 hours of shaking to ensure consistent tissue quality. Heavily-fixed human clinical samples were processed into FFPE blocks by following the same protocol used for FFPE embedding of mouse brain tissue.

#### FFPE block sectioning

In this study, FFPE blocks were sectioned at either 4 or 10 μm thickness. Blocks were first trimmed to a horizontal point using a Leica RM2255 microtome (Leica). The flattened blocks were sectioned at the desired thicknesses on silane-coated microslides (Muto). The sectioned slides were air-dried and stored in a cold room with silica gels to preserve sample quality.

#### H&E staining

Both 4 μm and 10 μm FFPE sections were used for hematoxylin and eosin (H&E) staining. H&E staining was performed using a Leica Autostainer XL, CV5030 (Leica) according to the following protocol: three changes of xylene for 7 minutes each, two changes of 100% ethanol for 1 minute each, two changes of 95% ethanol for 1 minute each, washed in water for 1 minute, stained in Harris hematoxylin solution for 5 minutes, washed in water for 1 minute, differentiated in 1% HCl for 3 seconds, washed in water for 30 seconds, blued in 0.3% ammonia solution for 3 seconds, counterstained in eosin Y for 20–60 seconds, rinsed twice in 95% alcohol for 30 seconds each, dehydrated three times in 100% ethanol for 1 minute each, cleared in three changes of xylene for 2 minutes each. All sectioning and staining protocols were performed at the Department of Histopathology, Avison Biomedical Research Center, Yonsei University.

#### FFPE sample homogenization, deparaffinization, and rehydration

FFPE blocks were removed from their embedding cassette and placed on 150 x 20 mm Petri dishes. External paraffin wax was removed manually with a razor blade, taking care not to remove any tissue. The shaved block was chopped into small pieces using a razor blade. Equal amounts of the minced tissue were placed in 2 mL tubes along with 5 mm stainless steel beads (Qiagen) and 1.8 mL of xylene (Duksan) or limonene (Sigma-Aldrich). Tissues were homogenized using the Tissue Lyser II device (Qiagen) at 30 Hz for 4 minutes in a chemical fume hood. The collected tissue solution was pelleted at 1,000 g for 3 minutes at room temperature. The supernatant was discarded, and the pellet was resuspended in xylene and incubated at room temperature for 5 minutes and repeated once. Deparaffinization and rehydration steps were performed by rotating the sample at 15 rpm at room temperature. The tissue was then resuspended in prechilled 100% ethanol and incubated at room temperature for 5 minutes and repeated once. This was followed by a single incubation step with 90% and 70% ethanol. Tissues were pelleted at 1,000 g for 3 minutes using a 4°C maintained bucket centrifuge from a 90% ethanol incubation step. Rehydrated tissue was pelleted at 1,000 g for 3 minutes using a 4°C maintained bucket centrifuge and resuspended in BCL solution (3X SSC, 3% PVSA, 0.025% Tween-20) and processed through a 20 μm cell strainer (tissues were first processed through a 100 μm cell strainer if necessary). Filtered tissue was washed twice more in BCL solution before additional crosslinking treatment.

For 4 or 10 μm FFPE sections, the mounted tissue was manually removed from the glass slide by scraping it off with a razor blade. Great care was taken to preserve the “scroll” of each sample. The collected scrolls of FFPE sections were then deparaffinized and rehydrated as described above for FFPE blocks. H&E-stained sections were also collected by manually scraping off the sample from the slides. H&E-stained sections were only rehydrated as they were already deparaffinized during the staining protocol. Sections were not further homogenized after rehydration. Before each experiment with FFPE or H&E sections, one to two slides were run through the FX-seq protocol to preliminarily assess the yield of nuclei and RNA quality.

#### Heavily fixed tissue homogenization

Similar to FFPE block homogenization, heavily PFA-fixed tissue without paraffin embedding was also divided and homogenized using a Tissue Lyser II device (Qiagen). However, chilled HM solution (10 mM Tris pH 7.0, 3% PVSA, 21 mM MgCl_2_, 1 mM CaCl_2_) was used instead of xylene or limonene for heavily PFA-fixed tissues. Note that the HM solution was modified from previously published methods for tissue dissociation by adding PVSA^47^. Homogenized tissue was collected and processed sequentially through 100 μm and then through a 20 μm cell strainer. Filtered tissue was washed for a total of three times by pulling down at 1,000 g for 3 minutes using a 4°C maintained bucket centrifuge in HM solution and immediately used for further downstream protocol or stored in a deep freezer until further use.

#### Fixative eXchange

Homogenized and washed tissue samples were first permeabilized in PBST3 solution (1X PBS, 0.2% TritonX-100, 3% PVSA) for 15 minutes at room temperature by rotating at 3 rpm. Samples were pelleted at 1,000 g for 3 minutes using a 4°C maintained bucket centrifuge and the pellet was collected and incubated in CL(–) solution (1X MOPS, 3% PVSA, 0.025% Tween-20) for 5 minutes in a cold room and repeated once. The CL(+) solution (1X MOPS, 3% PVSA, 0.025% Tween-20, 2X crosslinker) was added and shaken at 25°C for 15 minutes to crosslink the RNA molecules. After crosslinking, the tissue was pelleted at 1,000 g for 3 minutes using a 4°C maintained bucket centrifuge. The supernatant was discarded and the collected pellet was washed once in CT(–) solution (200 mM Tris pH 8.3, 3% PVSA). After centrifugation at 1,000 g for 3 minutes using a 4°C maintained bucket centrifuge, the pellet was carefully resuspended in CT(+) solution (200 mM Tris pH 8.3, 50 mM cat. 2, 3% PVSA, 8% w/v PEG-8000) and shaken at 55°C for 30 minutes to remove PFA crosslinking. Tissues after PFA removal were then pelleted at 1,000 g for 3 minutes using a 4°C maintained bucket centrifuge and the supernatant was discarded and proceeded further for nucleus isolation.

#### Nucleus isolation

The pelleted, FX-treated tissue was washed once with DG(–) solution (50 mM Tris pH 7.0, 10 mM EDTA pH 8.0, 10 mM NaCl, 1% PVSA, 0.2% Tween-20). The pellet was collected by pulling down at 1,000 g for 3 minutes using a 4°C maintained bucket centrifuge and resuspended in DG(+) solution (50 mM Tris pH 7.0, 10 mM EDTA pH 8.0, 10 mM NaCl, 1% PVSA, 0.2% Tween-20, 0.1 mg/mL Proteinase K) and shaken at 37°C for 20 minutes. Tissue digestion was optimized for each tissue type, as the ECM of some tissues and organs is more rigid and less susceptible to digestion. For instance, all human tissues analyzed in this study were digested with a DG(++) solution (50 mM Tris pH 7.0, 10 mM EDTA pH 8.0, 10 mM NaCl, 1% PVSA, 1% Tween-20, 0.1 mg/mL Proteinase K) and shaken at 37°C for 30 minutes. After digestion, the digested tissue was pelleted at 1,000 g for 3 minutes using a 4°C maintained bucket centrifuge. The supernatant was discarded, and the digested pellet was thoroughly resuspended in sucrose solution (1X PBS, 1% PVSA, 1.5 M sucrose) and centrifuged at 3,000 g for 20 minutes using a 4°C maintained bucket centrifuge to pellet the nuclei. For sectioned samples, the digested pellet was briefly filtered through a 40 μm cell strainer prior to sucrose gradient isolation. The supernatant was discarded with taking much care not to disturb the pellet. The pelleted nuclei were further permeabilized with PBST1 solution (1X PBS, 0.2% TritonX-100, 1% PVSA) for 15 minutes at room temperature by rotating at 3 rpm. Permeabilized nuclei were pelleted at 1,000 g for 3 minutes using a 4°C maintained bucket centrifuge. The second permeabilization step was omitted for sectioned samples because excessive TritonX-100 treatment resulted in severe RNA leakage from the physically sectioned nuclei. The collected nuclei were then washed for a total of three times in BR solution (5X SSC, 0.025% Tween-20, 0.1% v/v SUPERase-In) by pulling down at 1,000 g for 3 minutes using a 4°C maintained bucket centrifuge. After the final washing step, the nuclei were resuspended in PBSRI (1X PBS, 0.04% w/v BSA, 1% v/v SUPERase-In) (NOTE: PVSA inhibits reverse transcription and should not be used in this step). Nuclei were stained with DAPI (Invitrogen) and counted using an EVOS imaging system (Life Technologies).

#### Preparation of biological triplicates from PFA-fixed or FFPE mouse brains

For the validation of the FX-seq treatment in PFA-fixed or FFPE mouse brains, brain tissue was collected from three litter-mate mice. Each brain was processed separately but in parallel until the reverse transcription step. Control groups were not subjected to any additional crosslinking or organocatalyst treatment prior to enzymatic digestion. Tris control groups were not additionally crosslinked but treated at 55°C for 30 minutes in tris-only solution (200 mM Tris pH 8.3, 3% PVSA, 8% w/v PEG-8000). The Cat. 2 groups underwent organocatalyst treatment without additional crosslinking. The FX groups were additionally crosslinked and treated with organocatalysts prior to enzymatic digestion. Each brain of the biological triplicate was separated into 4 groups (No treatment control, Tris Ctrl, Cat. 2, FX) and given separate barcodes during reverse transcription and pooled during ligation and processed together in the subsequent steps of library construction. Each barcode was deconvoluted during computational analysis to distinguish between each experimental group and each brain.

#### Freshly extracted and lightly fixed versus heavily fixed mouse brain with FX-seq comparison experiment

To compare the performance of FX-seq with fresh samples, we followed the tissue preparation guidelines outlined in the Easy-sci-RNA-seq protocol^24^. Briefly, a fresh mouse brain was obtained after euthanasia and decapitation. Fresh brain tissue was minced into small pieces with physical exclusion of the cerebellum in cold buffer (1X PBS, 1% DEPC), and centrifuged at 200 g for 5 minutes. The 1 mL of EZ lysis buffer (Sigma-Aldrich) with 10 μL DEPC was added to the pellet and gently suspended 10 times with a wide-bore tip. The solution was placed on ice for 5 minutes and processed through a 40 μm cell strainer (Pluristrainer). An additional 500 μL of EZ lysis buffer with 0.1% SUPERase-In (Invitrogen) was added to the top of the strainer to process any remaining tissue. Processed tissue was centrifuged at 500 g for 5 minutes and resuspended in 1 mL of EZ lysis buffer with 0.1% SUPERase-In (Invitrogen) and pipetted three times. The solution was centrifuged at 500 g for 5 minutes and was briefly fixed in 0.1% PFA (EMS) and 1X PBS (Gibco) for 10 minutes on ice. Nuclei were pelleted at 500 g for 3 minutes and resuspended in 500 μL EZ lysis buffer with 0.1% v/v SUPERase-In (Invitrogen) and pipetted three times and repeated once more. The resulting nuclei were washed with 500 μL of wash solution (10mM Tris pH 7.5, 10mM NaCl, 3mM MgCl_2_, 0.01% Triton-X). Nuclei were resuspended in 500 μL of wash solution, briefly sonicated at the lowest setting for 12 seconds, and filtered through a 20 μm cell strainer. The cell strainer was washed with an additional 250 μL of wash solution. The sample was centrifuged at 500 g for 5 minutes and was resuspended in suspension solution (10mM Tris pH 7.5, 10mM NaCl, 3mM MgCl_2_). Nuclei were counted, resuspended in PBSRI, and reverse transcription was performed as described above. A paired litter-mate mouse was anesthetized, perfused, and shaken in 4% PFA (EMS), 1X PBS (Gibco) for 48 hours in a cold room in advance. The fixed litter-mate brain was treated with the FX-seq protocol as described above but with the resection of the cerebellum and given separate barcodes during reverse transcription. The separate barcodes were deconvoluted after the experiment to distinguish the transcriptomes of individual samples. Both techniques were performed in parallel.

#### FX-seq of human colorectal cancer FFPE sections with spatial contexts

For spatially contextualized FX-seq of human colorectal cancer (CRC) FFPE sections, an expert pathologist first annotated tumor and non-tumor regions by analyzing H&E sections. Based on these annotations, adjacent 10 μm FFPE sections were separated by physical scraping with a razor blade. The separated sections were simultaneously deparaffinized, rehydrated, and subjected to the FX-seq treatment. Detailed methods including the downstream protocol can be found above.

#### Reverse transcription & ligation

Combinatorially barcoded reverse transcription for sequencing experiments was performed according to the protocol described below. For each well, 2 μL of 5X Maxima H Minus reaction buffer (Thermo Fisher Scientific), 0.5 μL of 200U/μL Maxima H Minus reverse transcriptase (Thermo Fisher Scientific), 0.5 μL of 20U/μL SUPERase-In (Invitrogen), 2 μL of a mixture of 25 μM barcoded oligo-dT primer and 2.5 mM dNTP, 3 μL of 40% w/v PEG-8000 (Sigma-Aldrich), 2 μL of counted and prepared nuclei (7.5k–10k nuclei/μL) were thoroughly mixed and briefly pulse centrifuged using a 4°C maintained plate centrifuge. Each plate was incubated for 30 minutes at 55°C for reverse transcription. After reverse transcription, 30 μL of wash solution (1X PBS, 0.025% Tween-20) was added to each well, pooled into a single 5 mL tube, and centrifuged at 1,000 g for 3 minutes using a 4°C maintained bucket centrifuge. Centrifuged nuclei were resuspended in 270 μL nuclei suspension buffer (1X PBS, 0.04% w/v BSA, 1% v/v SUPERase-In) per 96-well plate. Resuspended nuclei were ligated according to the steps described following. For each well in the ligation plate, 0.5 μL quick ligase, 5μL of 5X quick ligation Buffer, 2 μL of 10 μM barcoded adapter, 2.5 μL of nucleus suspension solution were thoroughly mixed and briefly pulse centrifuged. Each plate was incubated for 10 minutes at 25°C. After ligation, 30 μL of wash solution (1X PBS, 0.025% Tween-20) was added to each well, pooled in a single 5 mL tube, and centrifuged at 1,000 g for 3 minutes. The pelleted nuclei were filtered with a Flowmi cell strainer (SP Bel-Art) and another round of the washing step was performed before counting the nuclei. Counted nuclei were resuspended in wash solution (1X PBS, 0.025% Tween-20) at 200-500 nuclei/μL, depending on the number of cells to be sequenced. The remaining nuclei were stored at –80°C until further use. All oligonucleotide barcode sequences used in this study are from sci-RNA-seq3^23^.

#### Library preparation and sequencing

An aliquot of 5 μL of resuspended nucleus solution after ligation was combined with 3 μL of elution buffer (Qiagen), 1.33 μL of second strand synthesis buffer (NEB), and 0.66 μL of second strand synthesis enzyme mix (NEB) per well. The resulting mixture was incubated for 1 hour at 16°C. For tagmentation, 4 μL of 5X tagmentation buffer (50 mM TAPS-NaOH pH 8.5, 25 mM MgCl_2_, 50% DMF), 5 μL nuclease-free water, 1 μL Tn5 (in-house preparation described below, concentration properly titrated during optimization for library construction of each sample type) was mixed on ice and incubated at 55°C for 7 minutes. For inactivation, 5 μL of 0.1% SDS (Sigma-Aldrich) was added to each well and thoroughly resuspended. Inactivation was performed at 55°C for 15 minutes. After inactivation, 1 volume of SPRIselect was added to each well and washed and eluted to 16.5 μL according to the manufacturer’s protocol. For PCR enrichment, 20 μL of 2X NEB next high-fidelity PCR master mix (NEB), 2 μL of 10 μM barcoded Illumina P5 adapter, 2 μL of 10 μM barcoded Illumina P7 adapter, 16 μL of the eluted library solution were thoroughly mixed. Enrichment was performed using the following cycle setting: 72°C for 5 minutes, 98°C for 30 seconds, 9–14 cycles of (98°C for 10 seconds, 66°C for 30 seconds, 72°C for 1 minute), and a final extension step at 72°C for 5 minutes. The amplified library was size-selected using 0.8X volume of SPRIselect. All barcoded adapter sequences used in this study are from sci-RNA-seq3^23^. The final library concentration was determined by Qubit 4.0 fluorophore (Invitrogen) using 1X dsDNA HS assay (Invitrogen). Final library quality was determined by Agilent 4200 Tapestation (Agilent) using HSD1000 screentape (Agilent). Libraries were sequenced by the NextSeq 2000 instrument (Illumina) using P2 (2 × 100 bp) or P3 (2 x 150 bp) paired-end kits (manufacturer’s specs: P2 ∼400M and P3 ∼1.2B reads; read 1: 34 cycle; read 2: 100 cycle). The detected cell numbers and reads per cell can be found in Table S15.

#### Synthesis of crosslinkers

Unless otherwise stated, the reactions were conducted under an argon or nitrogen atmosphere, in dry solvents. Commercially available reagents were used as received unless otherwise stated. Reaction temperatures were controlled by a heating mantle (Misung Scientific). C18 flash chromatography was conducted using CombiFlash Rf 150 purification system (Teledyne) and C18 Puchem flash column (pore size: 120Å, particle size: 50 µm, cartridge mass: 26 g, Purichem). Proton (^1^H) nuclear magnetic resonance (NMR) spectra were recorded on a Bruker Avance III HD 300 MHz instrument in MeOD-*d4* solution (Sigma Aldrich). Data for ^1^H NMR spectra are presented as follows: chemical shift (δ ppm) [multiplicity, coupling constant (Hz), integration]. Mass spectra were obtained using an InfinityLab LC/MSD with 1260 series LC.

Dichloro(ethylenediamine)platinum(II) [Pt(en)Cl_2_, 100 mg, 0.31 mmol, Sigma-Aldrich] and dimethylformamide (DMF, Sigma-Aldrich) were added to a round-bottom flask with a magnetic stir bar. Silver nitrate (AgNO_3_, 58 mg, 0.31 mmol, Sigma-Aldrich) in 6 mL of DMF was added dropwise, and the mixture was stirred in the dark for 16 hours at room temperature. The mixture was then filtered through a Millex-GV membrane filter (pore size: 0.22 *µ*m, Millipore) to remove the silver chloride (AgCl). An aliquot of NH_3_–PEG_11_–NH_3_ (65 mg, 0.12 mmol, Broadpharm) was added and stirred overnight at 50°C in the dark. After completion, the mixture was concentrated under reduced pressure. The residue was dissolved in 3 mL of Milli-Q water and stored at 4 °C overnight under dark condition. The mixture was filtered through a Millex-GV membrane filter (pore size: 0.22 *µ*m, Millipore) to remove insoluble yellow particles, [Pt(en)Cl(NO_3_)]. The filtered solution was concentrated under reduced pressure and purified by C18 flash column chromatography (0.1% formic acid in water and 0.1% formic acid in acetonitrile, with a gradient increase of acetonitrile from 0 to 100 % in 10 minutes). The solvent was removed from the isolated fractions under reduced pressure and the resulting residue was dissolved in 3 mL of MeOH. The solution was filtered through a Millex-GV membrane filter (pore size: 0.22 *µ*m, Millipore) for removal of remaining [Pt(en)Cl(NO_3_)], yielding a yellow oil (119 mg, yield: 88.2%, Figure S2A). ^1^H-NMR (300 MHz, MeOD-*d4*): d = 3.97–3.68 (m, 52H), 2.98–2.56 (m, 14H). LC-MS (ESI+) *m/z* (calculated for C_28_H_66_Cl_2_N_6_O_11_Pt_2_ [M–2H]^2+^: 562.68, found 562.4 (Figure S2B and C).

#### In-house Tn5 expression and purification

pTXB1-Tn5 was a gift from Rickard Sandberg (Addgene_60240). For the expression and purification of Tn5, we followed Sandberg’s method^48^ with slight modifications. The expression plasmid pTXB1-Tn5 was transformed into chemically competent cells (C3013, NEB) under ampicillin selection. To prepare 500 mL of the cell culture, 250 mL of LB medium (MP Biomedicals) was added to each of two 1 L flasks, and 5 mL of cells grown overnight were seeded into each 250 mL of medium. The cells were grown in LB medium (MP Biomedicals) with ampicillin at 37°C and shaken at 250 rpm until the optical density at 600 nm (A600) reached a range of 0.5 to 0.9. Then, the culture was cooled in a cold room for 30 minutes, and IPTG (SG bio) was added to a final concentration of 0.25 mM. After overnight growth at 250 rpm at 25°C, the culture reached an A600 of 3.0. The culture was then harvested and frozen at –80°C until further use.

For protein extraction and purification, the pellet from 500 mL of cultured cells was resuspended in 80 mL of HEGX buffer (20 mM HEPES-NaOH pH 7.2, 800 mM NaCl, 1 mM EDTA, 10% glycerol and 0.1% TritonX-100) supplemented with complete protease inhibitors (Roche). The subsequent sonication and poly(ethyleneimine) (PEI) precipitation steps were performed following Sandberg’s method. The clarified lysate was passed through a 0.45 μm filter (Millex), and 10 mL of chitin resin slurry (NEB) was added to the filtrate. The mixture was then transferred to an Econo-Pack chromatography column (Bio-Rad), and the resin was washed by gravity with 200 mL of HEGX buffer (20 times resin volume). The mixture was poured into two 50 mL Falcon tubes, and the tubes were rotated at 10 rpm in a cold room for one hour to allow the intein-tagged Tn5 to bind to the chitin resin. To remove the intein tags, we followed Sandberg’s method. An aliquot of 20 mL of elution buffer (HEGX buffer with 100 mM DTT) was added to the top of the column bed, and 5 mL of the buffer was allowed to flow through the column. The column was sealed for 48-72 hours at 4°C to allow cleavage of the inteins to release Tn5. Elution of Tn5 was performed in 3 mL aliquots. Each fraction was confirmed by SDS-PAGE, and the samples containing Tn5 were pooled and concentrated to 5 mL using an Amicon Ultra-15 centrifugal filter (Millipore). The sample was dialyzed overnight through two times of buffer exchanges of 1 L of 2X Tn5 dialysis buffer (100 mM HEPES-KOH at pH 7.2, 800 mM NaCl, 0.2 mM EDTA, 2 mM DTT, 10% glycerol). Note that we removed Triton X-100 and increased NaCl concentration to 800 mM in the dialysis buffers to facilitate long-term storage at –80°C^49^. Final protein concentration was measured using Pierce 660 nm protein assay reagent (Thermo Fisher). Tn5-MEDS B/B oligonucleotides were used for the formation of Tn5 transposome assembly according to Sandberg’s method. We obtained approximately 1 mg of purified Tn5 transposase in a single 500 mL culture, which is sufficient for 500-1,000 tagmentation reactions. The resulting Tn5 transposome can be stored at –80°C for long-term preservation and remains active after 2 years of storage.

#### Preprocessing of sequencing reads and generation of gene expression matrix

Fastq files generated by sequencing were processed following the analysis pipeline of sci-RNA-seq3^23^ with minor modifications. Briefly, generated reads were annotated by reverse transcription and ligation indexes within Levenshtein edit distance (ED) < 2. Individual reads were clipped using trim_galore v0.6.4 with AAAAAAAA sequence and default settings. Trimmed reads were mapped to the reference genome GRCm39 for mouse and GRCh38.p13 for human, using the default setting of STAR v.2.7.9a^50^ and gene annotations (GENCODE V42 for human and GENCODE VM27 for mouse). Duplicate reads were filtered using the unique molecular identifier (UMI) sequence, cell index with reverse transcription index, ligation adaptor index, and read 2 end-coordinate. Gene expression matrices of individual cells were generated using the HTSeq package^51^ by assigning mapped regions to intron- or exonic regions of each gene. In the case of multiple mapped reads, we assigned them to the closest end of the respective gene. However, if there were other genes within a distance of less than 100 bp from the closest gene, these reads were discarded.

#### Processing and visualization of single-nucleus RNA sequencing data

The gene expression matrix was constructed from the identified gene count during the preprocessing step using the Python package, scanpy v1.9.0^52^. Downstream analyses without detailed information on the analytical metric can be found in Table S16. Cells that did not meet specific quality control parameters, including thresholds for UMIs, detected gene numbers, and mitochondrial read fractions, were discarded from the downstream analysis. After merging the initial gene matrix, which distinguished between intron- and exonic regions within individual genes, only genes containing protein-coding sequences, pseudogenes, and long non-coding RNAs were used for further analysis. Finally, genes with no counts in the entire data set were excluded from the analysis. Suspected doublets were identified using Scrublet^53^ with different parameters and filtered with manual thresholds. The expression matrix of cells was normalized to the total number of mean UMIs of all cells, followed by log transformation with the addition of a pseudo count. The data were then scaled to have zero mean and unit variance. The dimensionally reduced data set by principal component analysis (PCA) was subjected to the neighborhood graph computation, and cells were clustered using the Leiden algorithm and embedded in the UMAP with optimal parameters for each experiment using functions implemented in scanpy. Genes specific to each cluster were identified using scanpy.tl.rank_genes_groups functions followed by filtering with experiment-specific parameters. Clusters were assigned to specific cells based on the identified marker genes according to molecular criteria from previous studies. For mouse brain cell type annotation, we referenced the single-cell and single-nucleus transcriptome atlas and marker genes from previous studies^24,54–56,26^. For cell type annotation in human clinical samples, we referenced the Human Protein Atlas to identify marker-gene-specific cell types in each tissue^57,58^. Clusters showing combinations of marker genes from other clusters were further filtered out as potential doublet populations. For sub-cluster analysis, unprocessed gene matrices corresponding to cells in each sub-cluster were reanalyzed from the gene filtering step. To construct the pathways between the identified clusters, we applied the partition-based graph abstraction (PAGA) algorithm^59^ after ordering the cells according to the calculated diffusion pseudotime^60^. For CNV inference from the gene expression matrix in Figure 5, we used inferCNV v1.3.3^41^ to infer CNV states from epithelial lineage transcriptomics. Raw gene expression matrices from cells above 200 UMIs were provided to inferCNV and processed. Average read counts per gene below 0.001 were filtered to account for low reads per cell in the matrix. The 3 states (neutral, loss, and gain) of CNV were inferred using HMM3 methods and analyzed at the sample level for Figure 5 and at the individual cell level for Figure S20.

## Notes

### Competing Interest Statement

C.H.S. is a co-founder and the CEO of Pixelas. Y.T.L. is a co-founder and the CTO of Pixelas. FX-seq is the subject of pending patent applications, including filings in South Korea and a PCT application.

