## Supplementary Information for "Fixative eXchange (FX)-seq: Scalable Single-nucleus RNA Sequencing Analysis of PFA-fixed or FFPE Tissue"

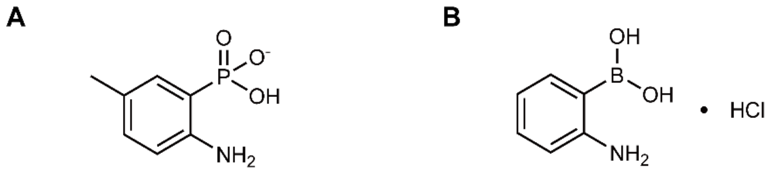


(2-Amino-5-methylphenyl)phosphonic acid
Cat. 1

2-Aminophenylboronic acid hydrochloride
Cat. 2

**Figure S1. Molecular structures of organocatalysts employed for FX-seq.**

(A and B) molecular structures of Cat. 1(A) and Cat. 2(B).

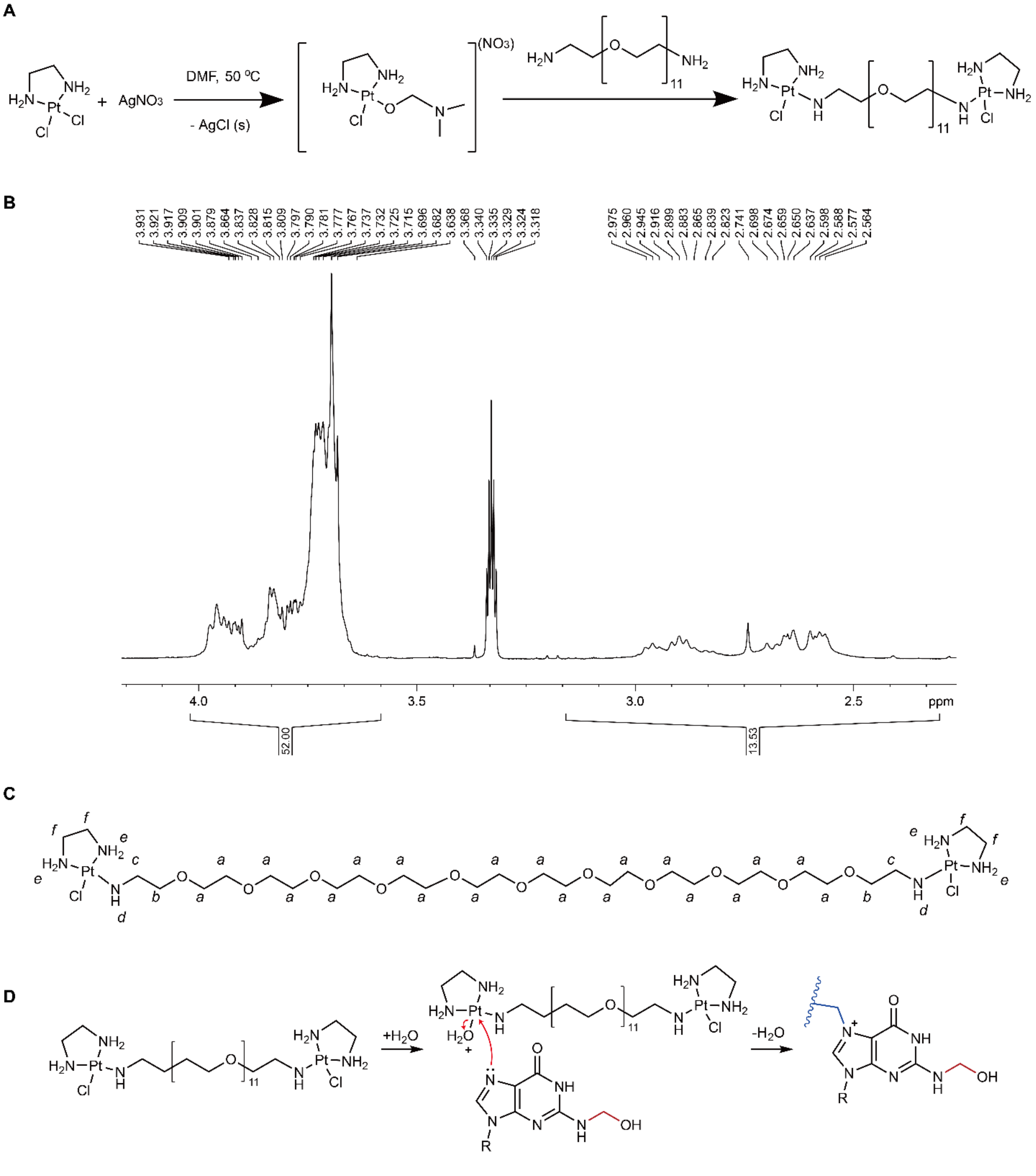


Mechanism of regiospecific reaction of Pt(II)-derivative cross-linker

**Figure S2. Synthesis and regiospecificity of the FX-seq crosslinker**

(A) Synthetic route for the crosslinker. Detailed steps are provided in the method section.

(B and C) $^{1}H$ NMR spectrum of the crosslinker (B) with (C) peak assignments.

(D) Guanine-N7 regioselective reaction of the crosslinker.

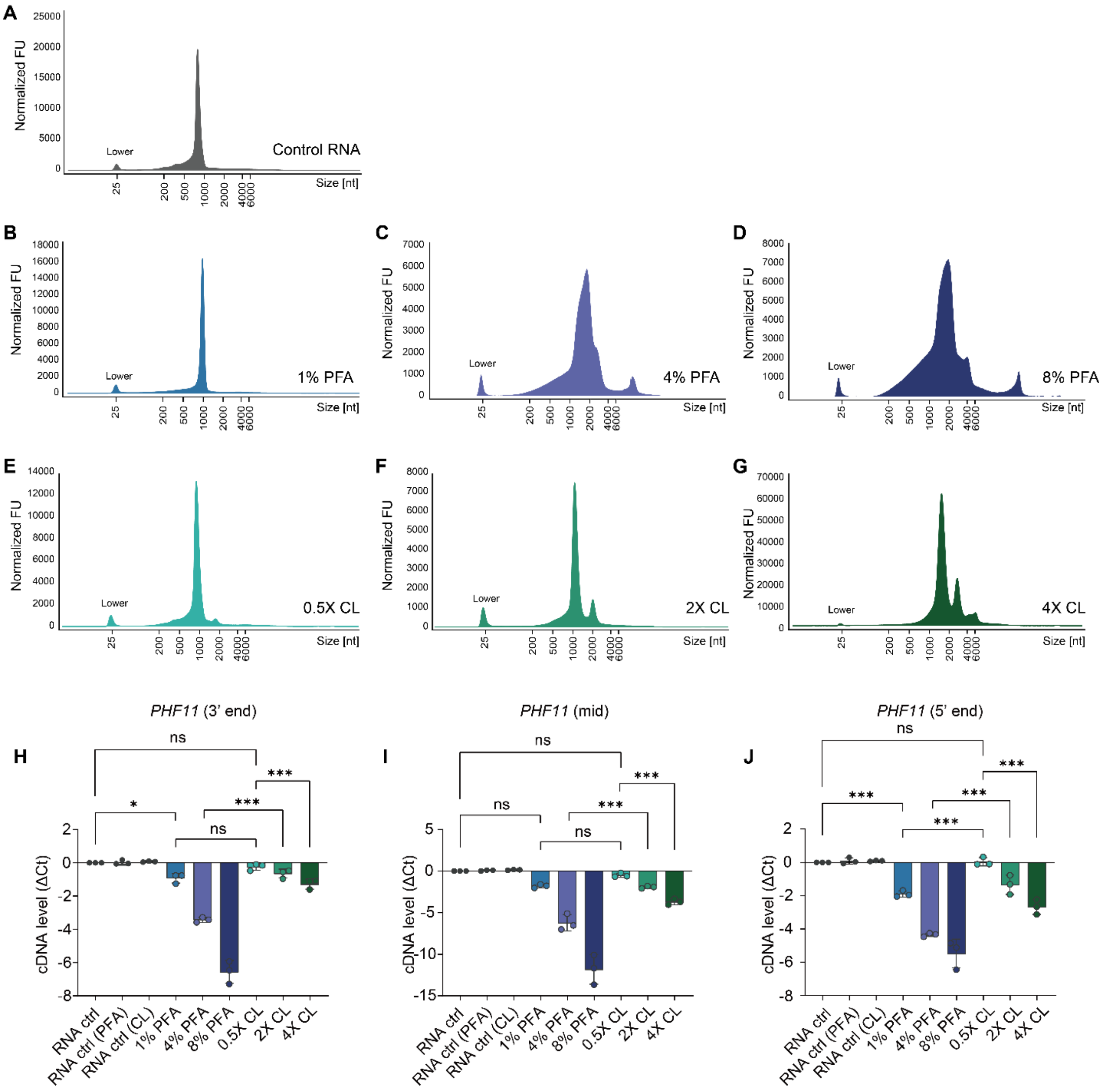


**Figure S3. Guanine-N7 regioselective crosslinker minimizes inhibition of *in vitro reverse transcription* (RT).**

(A–G) Automated gel electrophoresis data of crosslinked *in vitro* RNA species. Data are plotted by normalized fluorescence units (FU) per size (bp). "Lower" indicates low-molecular-weight species markers. Experimental conditions: (A) non-crosslinked RNA control, (B) 1% PFA, (C) 4% PFA, (D) 8% PFA, (E) 0.5X crosslinker (CL), (F) 2X CL, (G) 4X CL. Note that 4% PFA and 2X CL concentrations are the standard concentrations employed for tissue fixation.

(H–J) Bar graph showing relative inhibition by reactions with PFA or CL at corresponding concentrations. Due to difficulties in RNA purification, crosslinked RNAs from each condition were appropriately diluted to avoid inhibition of RT and qPCR reactions. RNA ctrl (PFA) and RNA ctrl (CL) groups contained parallel amounts of the respectively diluted crosslinkers to ensure that RT reactions were performed without crosslinking bias. ΔCt values of each experimental group were calculated by normalization with respect to the control group. Three different primer sets were designed to estimate the processivity of RT enzymes in progressive regions of the RNA species. Specifically, the primers targeted RNA from (H) 3' end, (I) middle region, and (J) 5' end. Note that the crosslinking effect of CL shows marginal inhibition whereas the crosslinking effect of PFA fixation shows significant and direct inhibition. We also found that the inhibitory effect of CL reactions appears to accumulate with increasing reach towards the 5' end, which requires longer processivity by RT enzymes. However, FX-seq is designed with 3' sequencing as the primary goal. Therefore, we expect the inhibitory effect of CL to outweigh the modest decrease in cDNA synthesis yield at full processivity. Data are represented as mean ± SD. *P* values (NS ≥ 0.05, **P* ≤ 0.05 ***P* ≤ 0.01 ****P* ≤ 0.001) were determined by unpaired one-way ANOVA followed by Bonferroni's multiple comparison test.

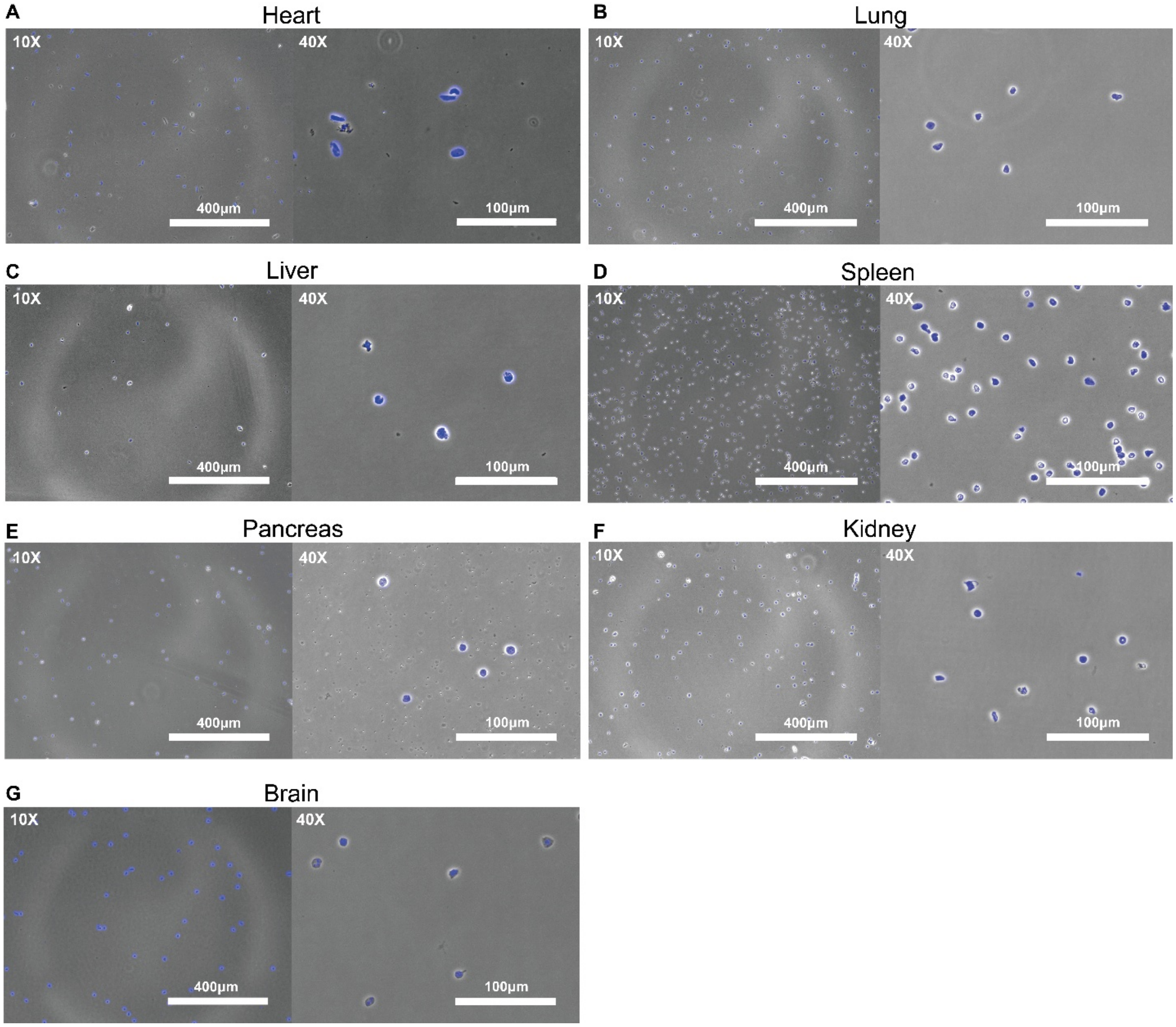


**Figure S4. Versatility of the FX-seq digestion strategy across multiple mouse organs.**

(A–G) Brightfield images of isolated single nuclei from perfused and heavily PFA-fixed mouse organs: (A) heart, (B) lung, (C) liver, (D) spleen, (E) pancreas, (F) kidney, and (G) brain. A detailed protocol is available in STAR Methods. Grayscale: phase contrast, blue: DAPI.

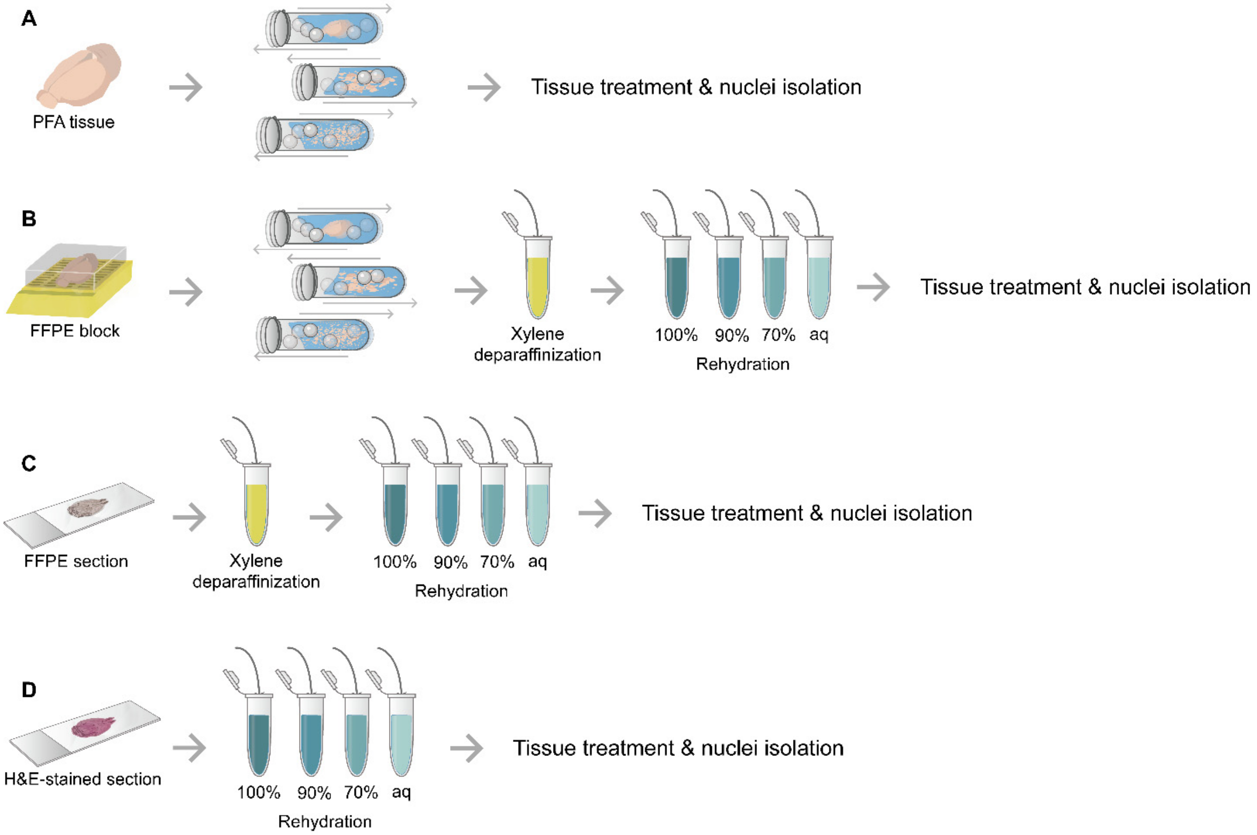


**Figure S5. FX-seq tissue preparation**

(A–D) Schematic diagram of the nucleus isolation strategy for PFA tissue (A), FFPE block (B), FFPE section (C), and H&E-stained section (D). Detailed protocols are available in STAR Methods.

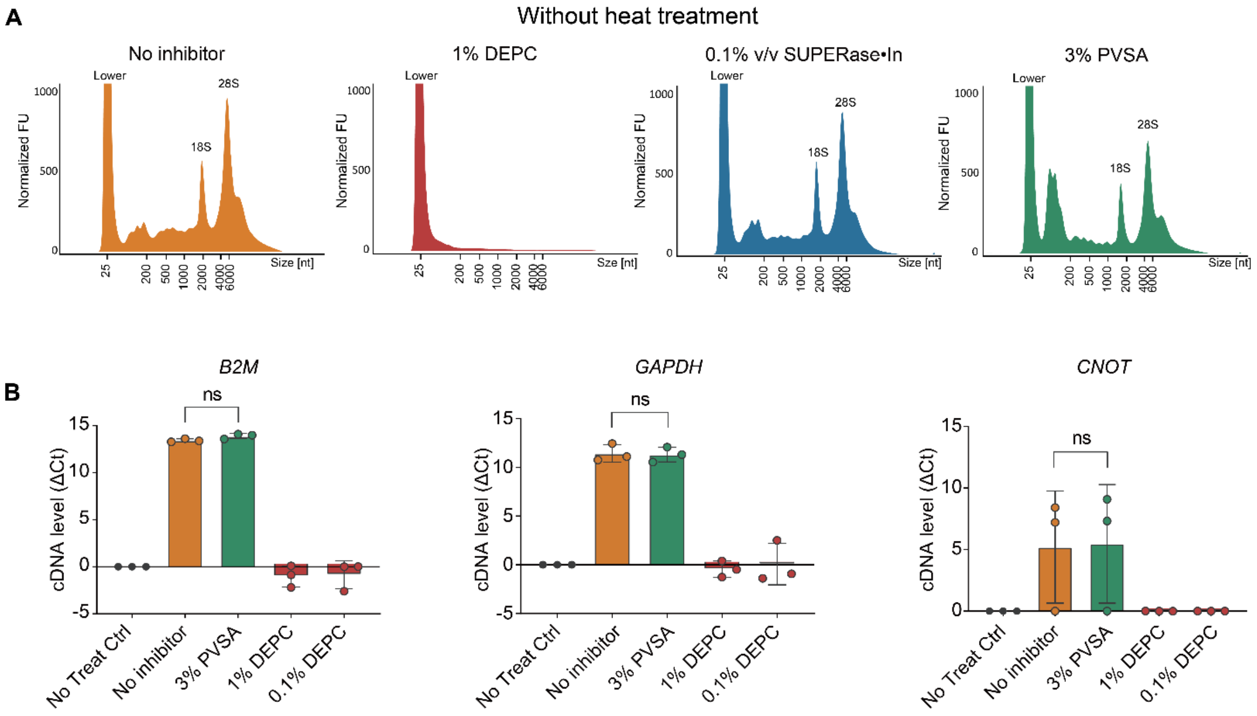


**Figure S6. PVSA serves as an effective, thermostable RNase inhibitor with enzymatic reaction compatibility for both the HeLa cell line model and heavily PFA-fixed mouse brain tissue.**

(A) Automated gel electrophoresis of total RNA extracted directly from heavily fixed mouse brain tissue without any heat treatment. Data are plotted by normalized fluorescence units (FU) per size (bp). "Lower" indicates low-molecular-weight species markers. No inhibitor (orange), 1% DEPC (red), 0.1% v/v SUPERase-In (blue), 3% PVSA (green) were employed as RNase inhibitors. Note that moderate heat treatment enhanced RNase activity, resulting in severe total RNA degradation in all other groups excluding 3% PVSA as shown in Figure 1G. Therefore, we incorporated PVSA into our protocol to serve as an inexpensive, thermally stable, and enzymatic reaction compatible RNase inhibitor.

(B) Comparison of RNase inhibitors in the HeLa cell lines confirmed by cDNA synthesis yield quantified by qPCR. The ΔCt values of each experimental group were calculated by normalization with respect to the control group. Three sets of genes were investigated: *B2M* (left), *GAPDH* (middle), *CNOT* (right). All three genes showed consistent results with the efficacy of PVSA in the FX-seq protocol. It should be carefully noted that while DEPC is an effective reagent that covalently blocks RNase enzymes, it also reduces all other enzymes to no activity. Since FX-seq employs enzymatic digestion, we conclude that DEPC is an incompatible RNase inhibitor here. The cDNA levels from each experimental group were normalized with respect to the no treatment control. Data are represented as mean ± SD. *P* values (NS ≥ 0.05, **P* ≤ 0.05 ***P* ≤ 0.01 ****P* ≤ 0.001) were determined by unpaired one-way ANOVA followed by Bonferroni's multiple comparison test.

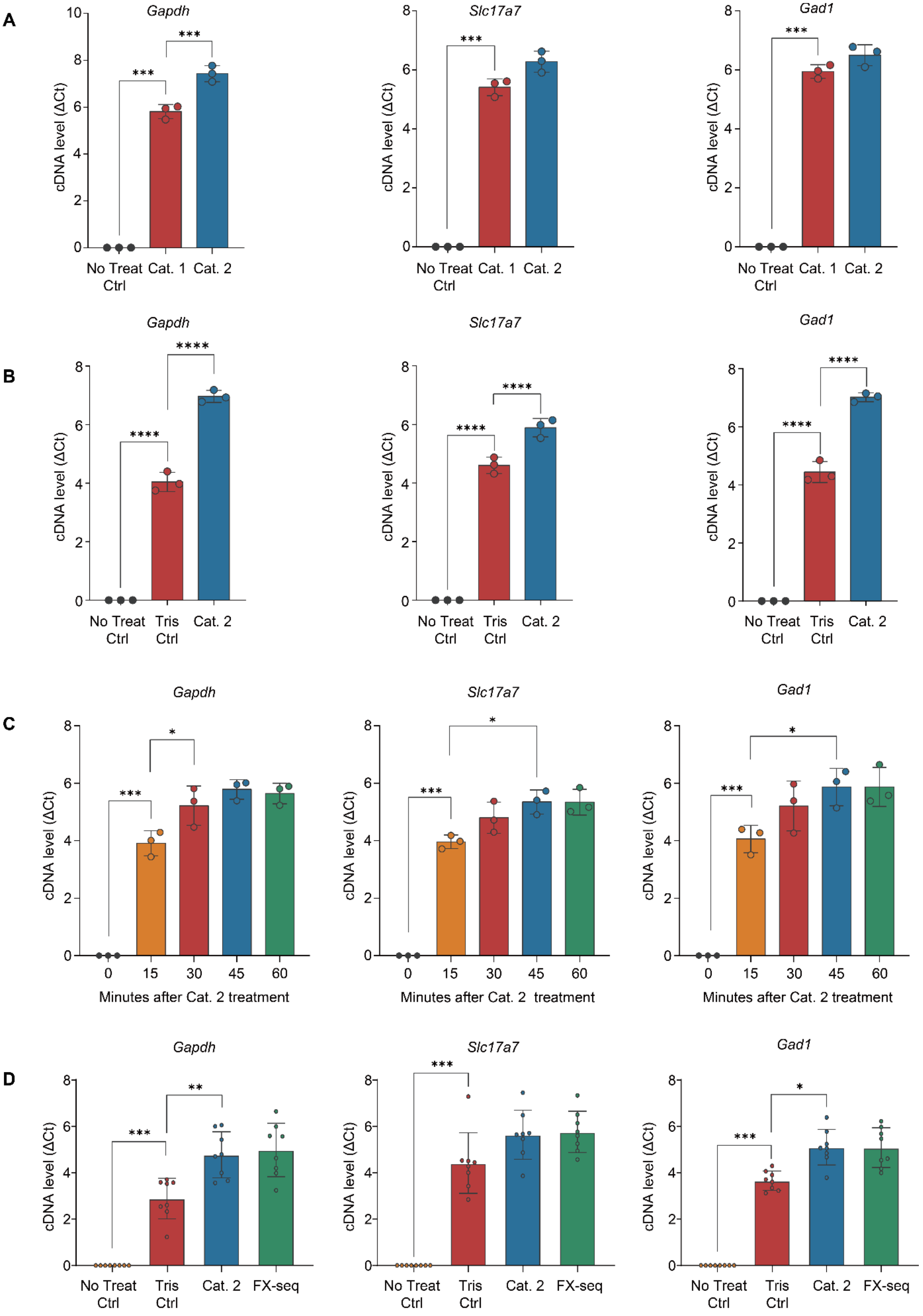
A
B
C
D
Gapdh
Slc17a7
Gad1
cDNA level (ΔCt)
No Treat Ctrl
Cat. 1
Cat. 2
Tris Ctrl
FX-seq
Minutes after Cat. 2 treatment
0
15
30
45
60
***
****
**
*

**Figure S7. *In situ* validation of the FX-seq catalyst in heavily PFA-fixed mouse brain tissue.**

Quantified cDNA levels in perfused and heavily fixed mouse brain tissue from qPCR assays. The ΔCt values of each experimental group were calculated by normalization with respect to the control group. Commonly enriched genes in the mouse brain were targeted: *Gapdh* (left), *Slc17a7* (middle) and *Gad1* (right).

(A) Comparison of reaction kinetics between cat. 1 and cat. 2 shows that cat. 2 has a more favorable kinetics compared to cat. 1.

(B) Comparison between Tris-buffer moderate heat treatment and cat. 2 treatment. Although moderate heat treatment in high-molar Tris buffer still increases cDNA synthesis yield in heavily PFA-fixed mouse brain tissue, the addition of cat. 2. further increases reverse transcription yields.

(C) Time-dependent evaluation of cat. 2 treatment. Prolonged treatment with cat. 2 shows a plateau in cDNA synthesis yield, most likely due to RNA molecule leakage, as previously confirmed in the HeLa cell line model.

(D) Stepwise evaluation of FX components. Moderate heating in Tris buffer (Tris ctrl), the addition of cat. 2 along with moderate heating in Tris buffer (cat. 2) and additional crosslinking treatment (FX-seq) are evaluated. Note that the same experimental groups were sequenced and analyzed as shown in Figure 1.

Data are presented as mean ± SD. $P$ values (NS ≥ 0.05, *$P \leq 0.05$ **$P \leq 0.01$ ***$P \leq 0.001$) were determined by unpaired one-way ANOVA followed by Bonferroni’s multiple comparison test.

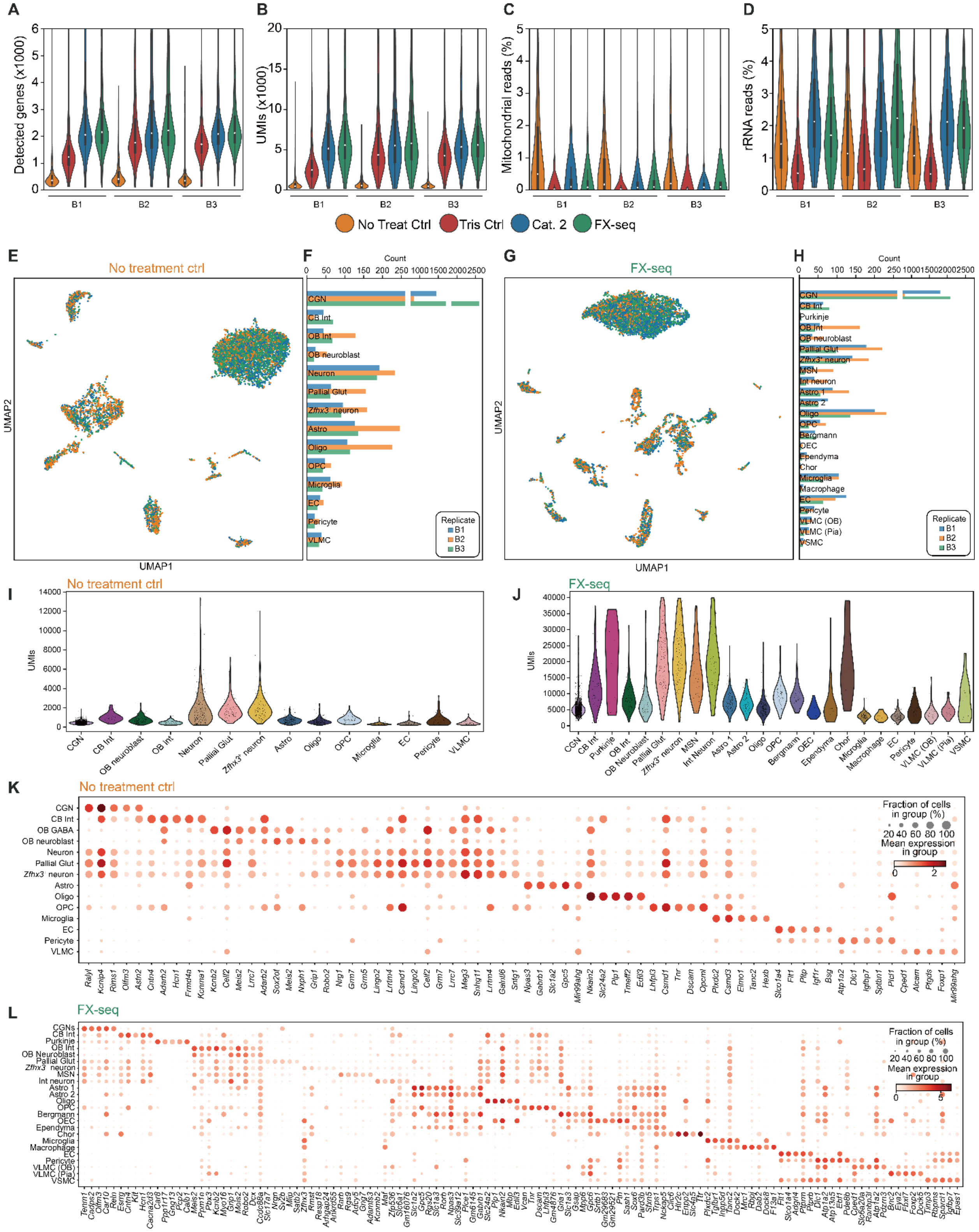


**Figure S8. Individual data from triplicate experiment of FX-seq using 4% PFA-fixed mouse brain.**

(A–D) Individual data of QC metrics from snRNA-seq of four different conditions: isolated nuclei without any treatment (No Treat Ctrl), moderate heat treatment with 200 mM Tris buffer (Tris Ctrl), moderate heat treatment with Tris buffered catalyst (Cat. 2), and the complete FX-seq procedure with additional crosslinking and tris buffered cat. 2 moderate heating (FX-seq). (A) Number of detected genes (A), number of UMIs per nucleus (B), ratio of mapped reads to mitochondrial genes (C), and ratio of mapped reads to rRNAs (D). All metrics showed similar values across biological replicates.

(E and F) Labelled nuclei from individual replicate from the UMAP analysis for no treatment control (E), and labelled cell count distribution across annotated clusters (F).

(G and H) Labelled nuclei from individual replicate from the UMAP analysis for FX-seq (G), and labelled cell count distribution across annotated clusters (H).

(I and J) Distribution of UMIs according to identified cell types in no treatment control (I), and FX-seq (J) in Figure 1.

(K and L) Dot plot of marker genes specific to identified cell types in no treatment control (K), and FX-seq (L) in Figure 1.

Abbreviations are as follows: CGN, cerebellar granule neuron; CB Int, cerebellar interneuron; OB Int, olfactory bulb interneuron; OB neuroblast, olfactory bulb neuroblast; Pallial Glut, pallial glutamatergic neuron; Astro, astrocyte; Oligo, oligodendrocyte; OPC, oligodendrocyte precursor; EC, endothelial cell; VLMC, vascular leptomeningeal cell; Purkinje, Punjinje cell; MSN, medium spiny neuron; Int neuron, interneuron; Bergmann, Bergmann glia; OEC, olfactory ensheathing cell; Ependyna, ependymal cell; Chor, choroid plexus epithelial cell; VLMC (OB), vascular leptomeningeal cell in olfactory bulb; VLMC (Pia), vascular leptomeningeal cell in pia mater; VSMC, vascular smooth muscle cell.

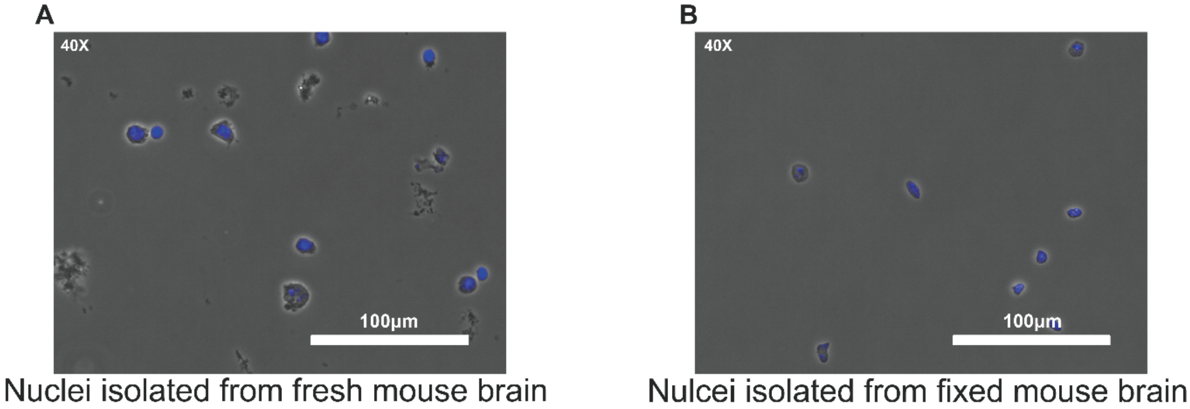


**Figure S9. Microscopic images of isolated nuclei from freshly extracted lightly PFA-fixed mouse brain, and heavily PFA-fixed mouse brain tissue.**

(A and B) Brightfield images of isolated nuclei in fresh mouse brain tissue (A), and perfused, heavily PFA-fixed mouse brain tissue (B). Grayscale: phase contrast, blue: DAPI.

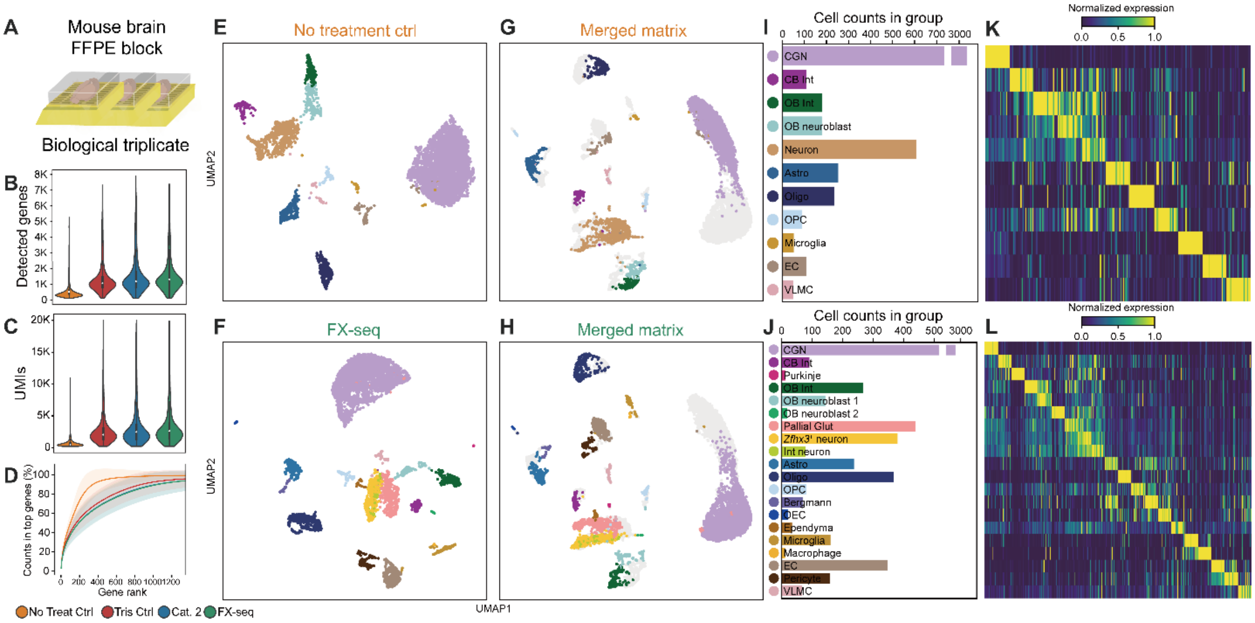

**Figure S10. Validation of FX-seq components in FFPE mouse brain tissue model.**

(A) Three litter-mate mouse brains were extracted immediately after euthanasia, fixed with PFA, and embedded in a paraffin block. Nuclei isolated from the FFPE block were subjected to RNA sequencing using FX-seq. Four different conditions were tested: isolated nuclei without any treatment (No Treat Ctrl), moderate heat treatment with 200 mM Tris buffer (Tris Ctrl), moderate heat treatment with Tris buffered catalyst (Cat. 2), and the complete FX-seq procedure with additional crosslinking and tris buffered cat. 2 moderate heating (FX-seq).

(B–D) QC metrics for FX-seq of FFPE mouse brain blocks. The individual components of FX-seq led to an increase in the number of detected genes (B), UMIs (C), and gene diversity (D). The optimized nucleus isolation protocol allowed the identification of various clusters even in the FFPE control samples, but the implementation of FX-seq significantly improved the resolution of cell type classification.

(E–L). UMAP clustering and cell-type annotation result of FFPE control samples (n = 4,914), and FX-seq (n = 5,852) after doublet removal. Individual UMAP visualization and annotation (E and F) and UMAP analysis of merged gene expression matrix (G and H) demonstrated improved analytical resolution of cell-type classification after FX-seq. Population distribution (I and J) and specificity of marker genes is visualized (K and L).

Abbreviations are as follows: CGN, cerebellar granule neuron; CB Int, cerebellar interneuron; OB Int, olfactory bulb interneuron; OB neuroblast, olfactory bulb neuroblast; Astro, astrocyte; Oligo, oligodendrocyte; OPC, olfactory bulb precursor; EC, endothelial cell; VLMC, vascular leptomeningeal cell; Purkinje, Punjinje cell; Pallial Glut, pallial glutamatergic neuron; Int neuron, interneuron; Bergmann, Bergmann glia; OEC, olfactory ensheathing cell; Ependyma, ependymal cell.

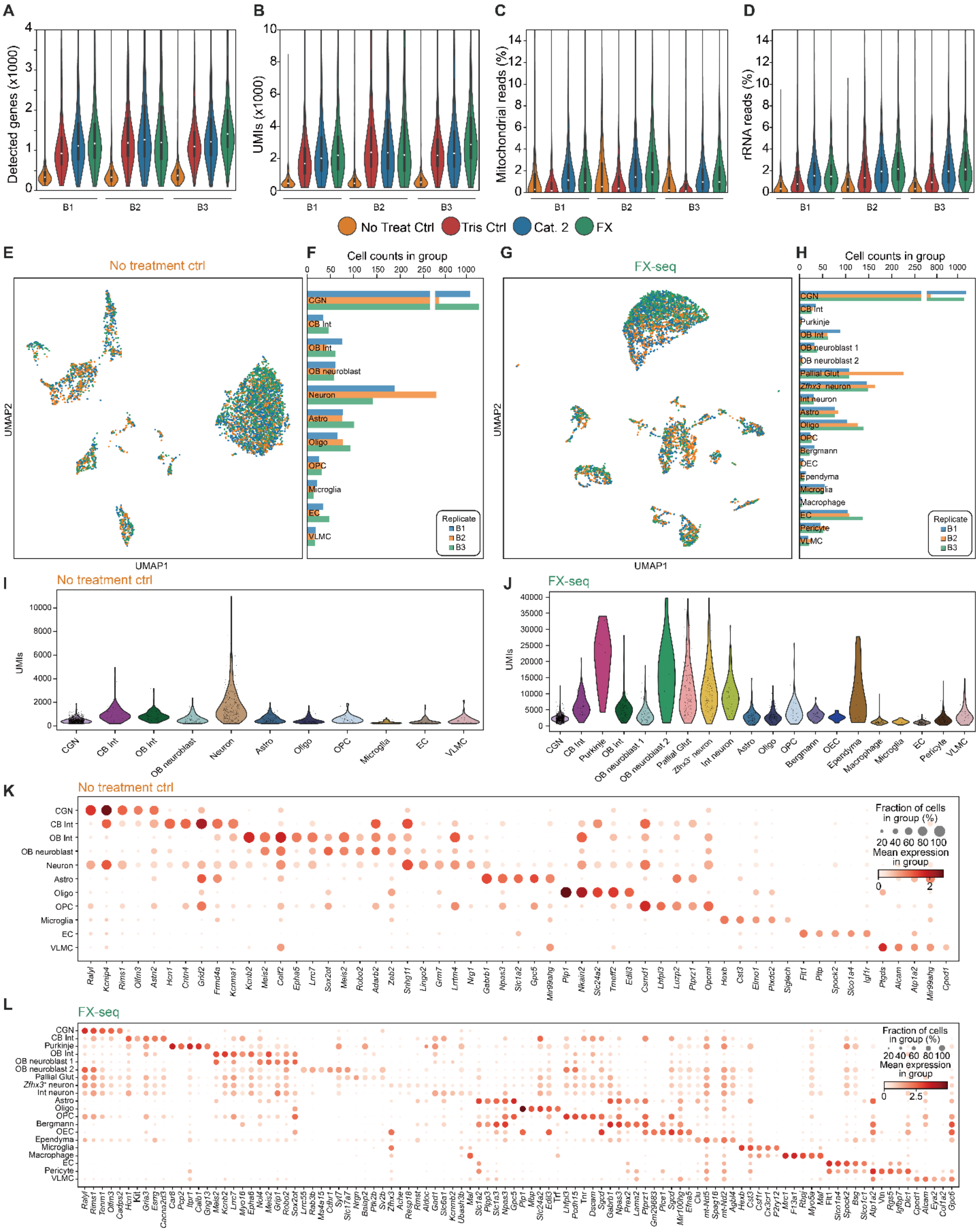


**Figure S11. Individual data from triplicate experiment of FX-seq using FFPE mouse brain blocks.**

(A–D) Individual data of QC metrics from snRNA-seq of four different conditions: isolated nuclei without any treatment (No Treat Ctrl), moderate heat treatment with 200 mM Tris buffer (Tris Ctrl), moderate heat treatment with Tris buffered catalyst (Cat. 2), and the complete FX-seq procedure with additional crosslinking and tris buffered cat. 2 moderate heating (FX-seq). Number of detected genes (A), number of UMIs per nucleus (B), ratio of mapped reads to mitochondrial genes (C), and ratio of mapped reads to rRNAs (D). All metrics showed similar values across the biological replicates.

(E and F) Labelled nuclei from individual replicate from the UMAP analysis for No treatment control (E), and labelled cell count distribution across annotated clusters (F).

(G and H) Labelled nuclei from individual replicate from the UMAP analysis for FX-seq (G), and labelled cell count distribution across annotated clusters (H).

(I and J) Distribution of UMIs according to identified cell types in no treatment control (I), and FX-seq (J) in Figure S9.

(K and L) Dot plot of marker genes specific to identified cell types in no treatment control (K), and FX-seq (L) in Figure S9.

Abbreviations are as follows: CGN, cerebellar granule neuron; CB Int, cerebellar interneuron; OB Int, olfactory bulb interneuron; OB neuroblast, olfactory bulb neuroblast; Astro, astrocyte; Oligo, oligodendrocyte; OPC, olfactory bulb precursor; EC, endothelial cell; VLMC, vascular leptomeningeal cell; Purkinje, Punjinje cell; Pallial Glut, pallial glutamatergic neuron; Int neuron, interneuron; Bergmann, Bergmann glia; OEC, olfactory ensheathing cell; Ependyma, ependymal cell.

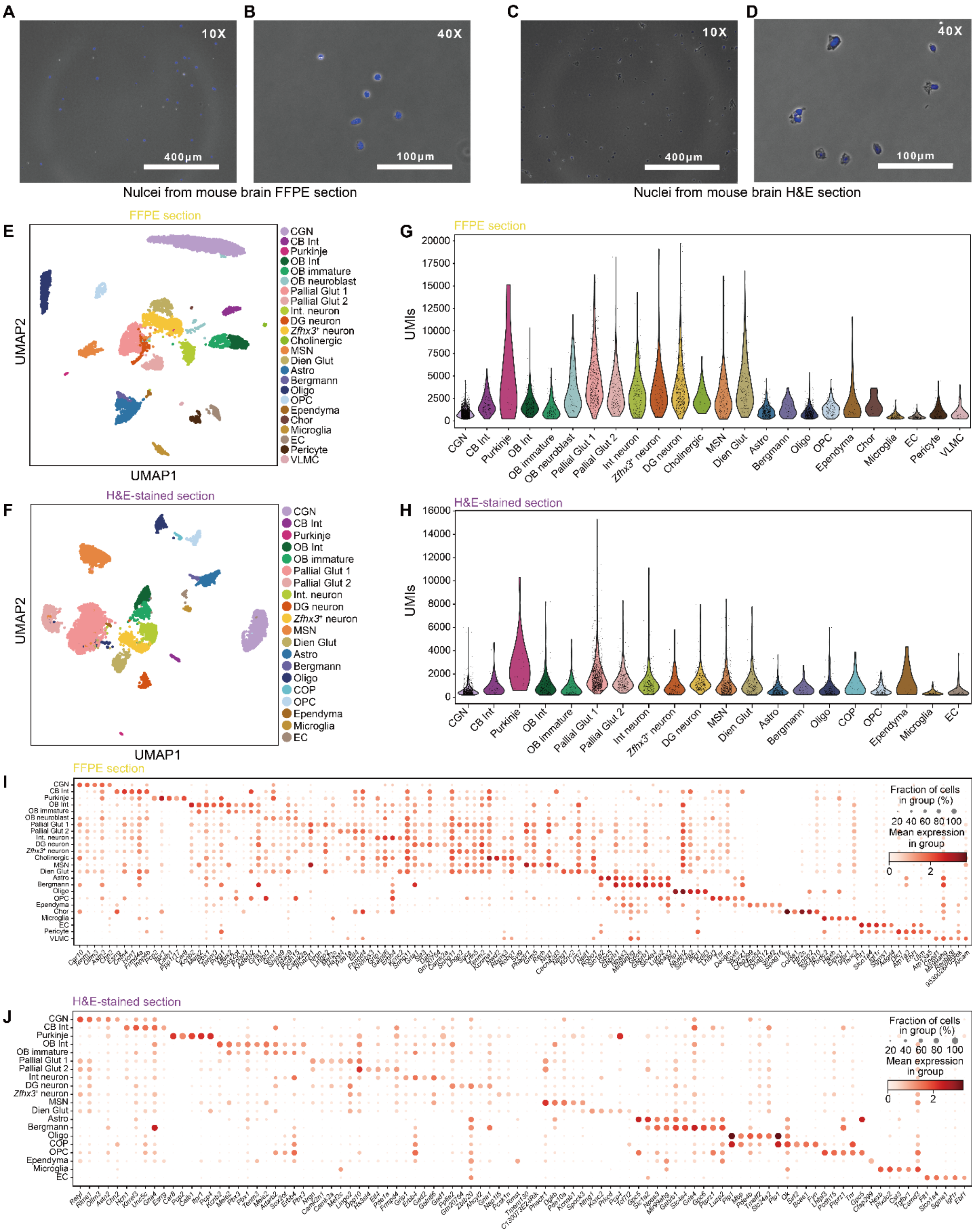


**Figure S12. Analysis result of FX-seq with mouse brain FFPE and H&E sections.**

(A and B) Microscopic images of mouse brain FFPE sections at (A) 10X magnification and (B) 40X magnification. Grayscale: phase contrast, blue: DAPI.

(C and D) Microscopic images of mouse brain H&E sections at (A) 10X magnification and (B) 40X magnification. Grayscale: phase contrast, blue: DAPI.

(E and F) The individually clustered UMAP analysis of mouse brain FFPE sections (E) and mouse brain H&E sections (F).

(G and H) UMI counts per annotated cell type in mouse brain FFPE sections (G), and mouse brain H&E sections (H) in Figure 3.

(I and J) DEG analysis showing marker genes in mouse brain FFPE sections (I) and mouse brain H&E sections (J) in Figure 3.

Abbreviations are as follows: CGN, cerebellar granule neuron; CB Int, cerebellar interneuron; Purkinje, Punjinje cells; OB Int, olfactory bulb interneuron; OB immature, olfactory bulb immature neuron; OB neuroblast, olfactory bulb neuroblast; Pallial Glut, pallial glutamatergic neuron; Int neuron, interneuron; DG neuron, dentate gyrus neuron; Cholinergic, habenula cholinergic neuron; MSN, medium spiny neuron; Dien Glut, diencephalic glutamatergic neuron; Astro, astrocyte; Bergmann, Bergmann glia; Oligo, oligodendrocyte; Ependyma, ependymal cell; Chor, choroid plexus epithelial cell; EC, endothelial cell; VLMC, vascular leptomeningeal cell; COP, committed oligodendrocyte precursor; OPC, oligodendrocyte precursor.

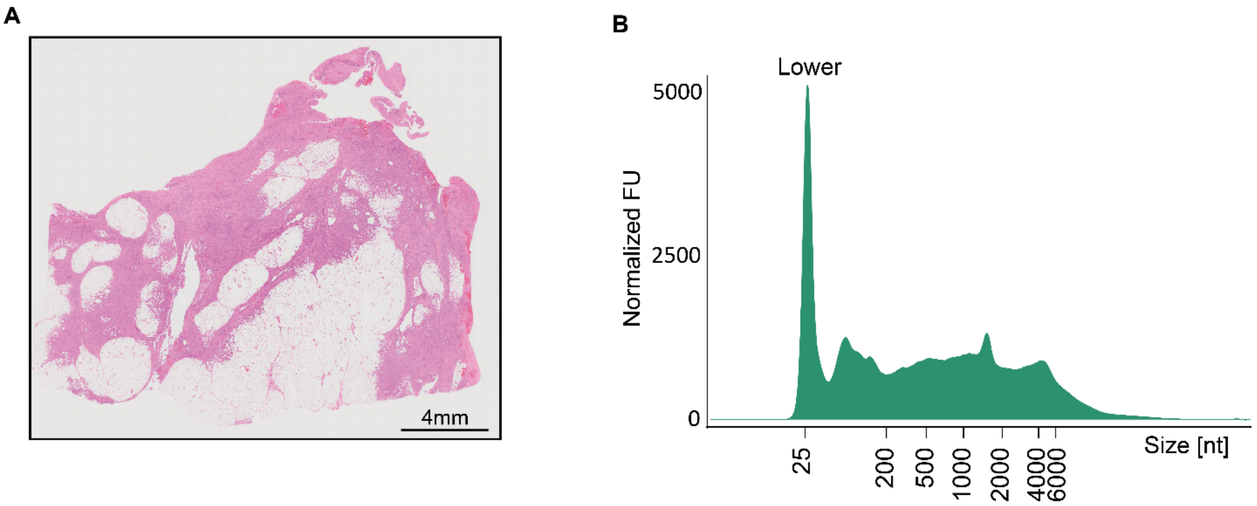


**Figure S13. Morphological and RNA quality assessment of omentum metastatic cancer.** (A) Microscopic image of 4 μm H&E-stained section of omentum metastatic cancer FFPE sample.

(B) Automated gel electrophoresis electropherogram of total RNA extracted from omentum FFPE sample. Data are plotted by normalized fluorescence units (FU) per size (bp). “Lower” indicates low-molecular-weight species markers.

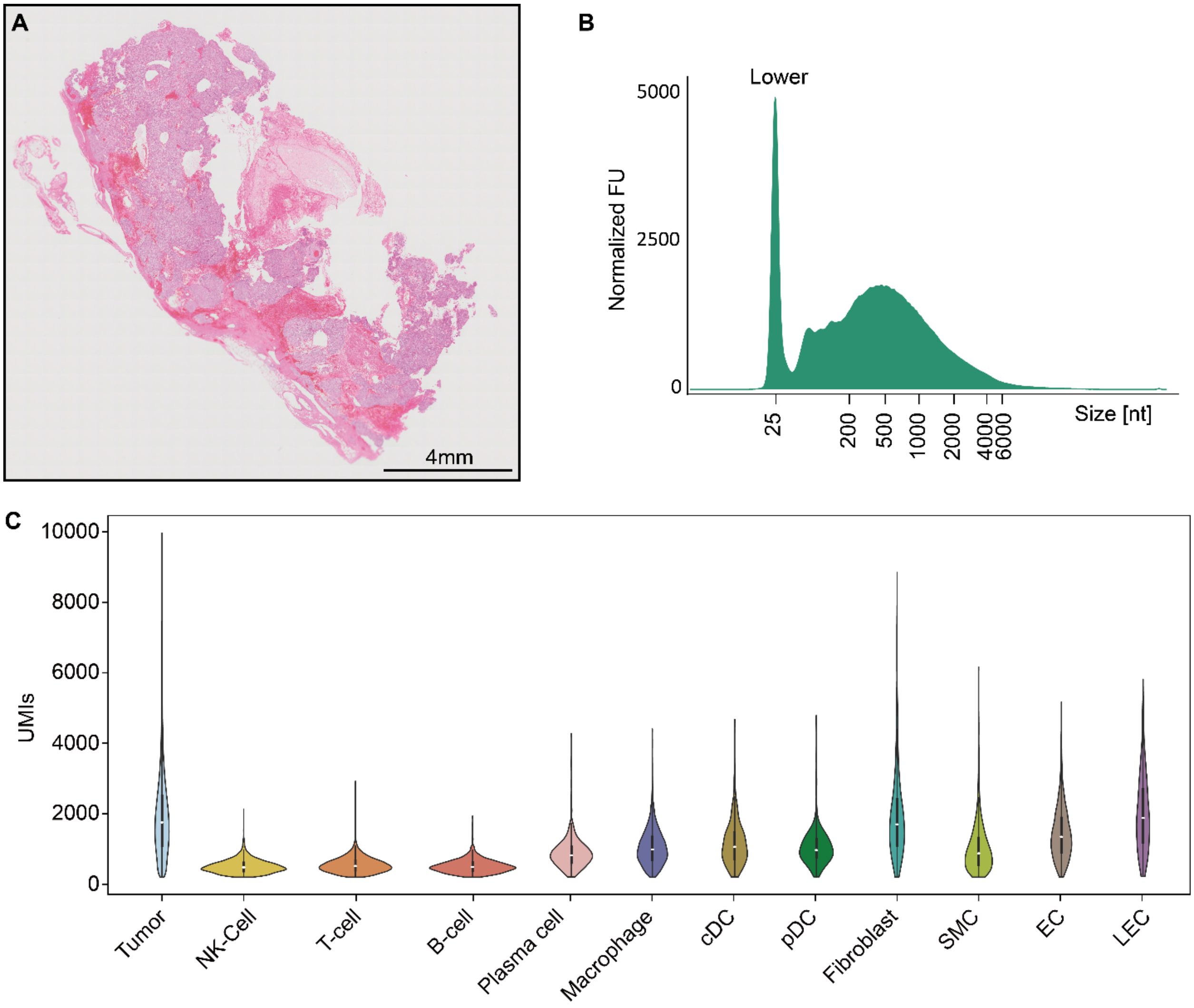


**Figure S14. Morphological and RNA quality assessment of gastrointestinal stromal tumor (GIST).**

(A) Microscopic image of 4µm H&E-stained section of GIST FFPE sample.

(B) Automated gel electrophoresis electropherogram of total RNA extracted from GIST FFPE sample. Data are plotted by normalized fluorescence units (FU) per size (bp). “Lower” indicates low-molecular-weight species markers.

(C) UMIs of annotated cell types shown in Figure 4.

Abbreviations are as follows: NK cell, natural killer cell; PC, plasma cell; MC, macrophage; cDC, classical dendritic cell; pDC, plasmacytoid dendritic cell; FB, fibroblast; SMC, smooth muscle cell; EC, endothelial cell; LEC, lymphatic endothelial cell.

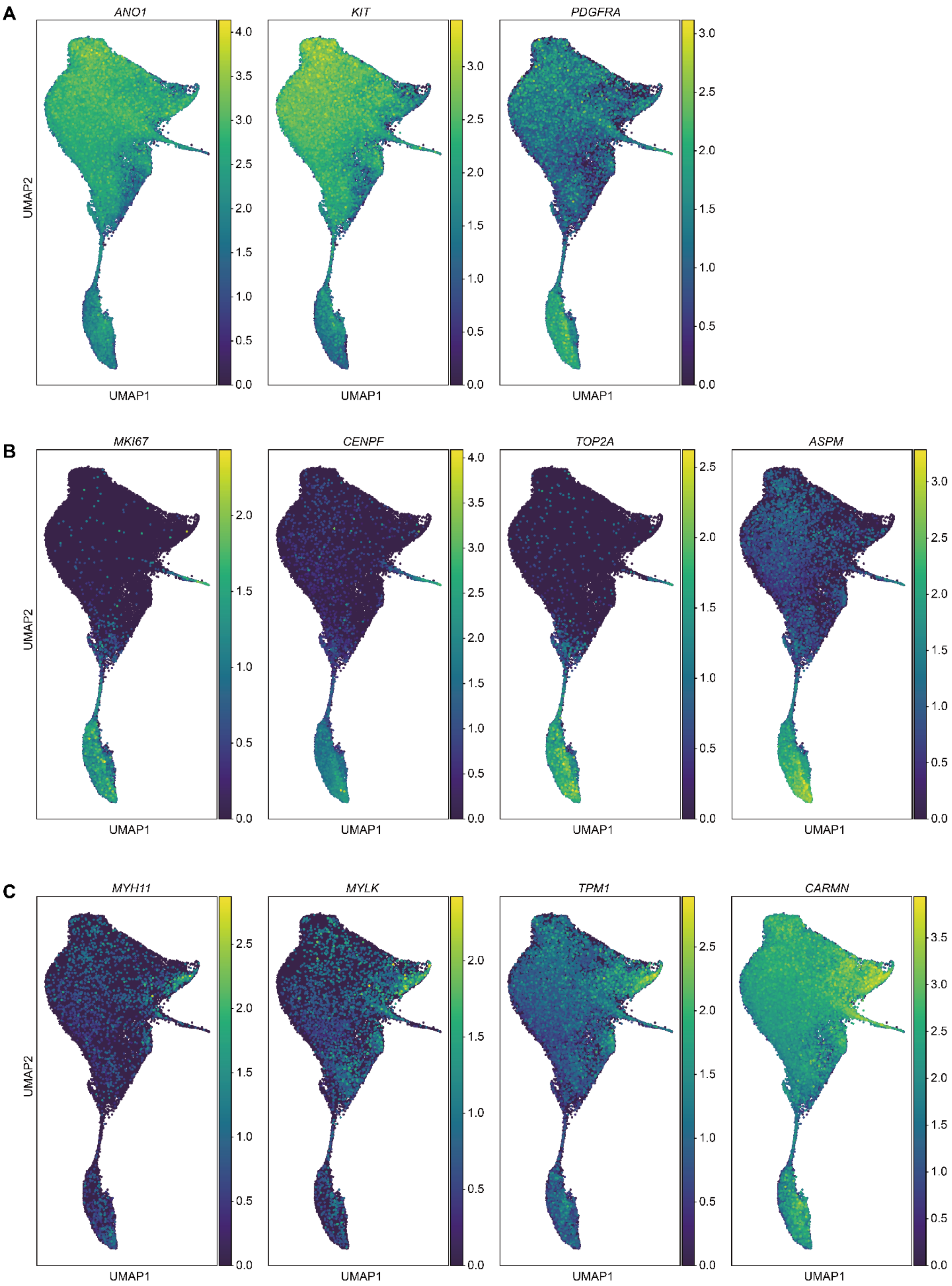
A
ANO1
KIT
PDGFRA
UMAP2
UMAP1
B
MKI67
CENPF
TOP2A
ASPM
C
MYH11
MYLK
TPM1
CARMN

**Figure S15. Differential gene expression profiles of tumor population in gastrointestinal tumor (GIST).**

(A) Expression of GIST marker gene (*ANO1*) and GIST oncogenic drivers (*KIT* and *PDGFRA*).

(B) Expression of proliferation marker genes.

(C) Expression of genes related to the functionality of interstitial cells of Cajal (ICC).

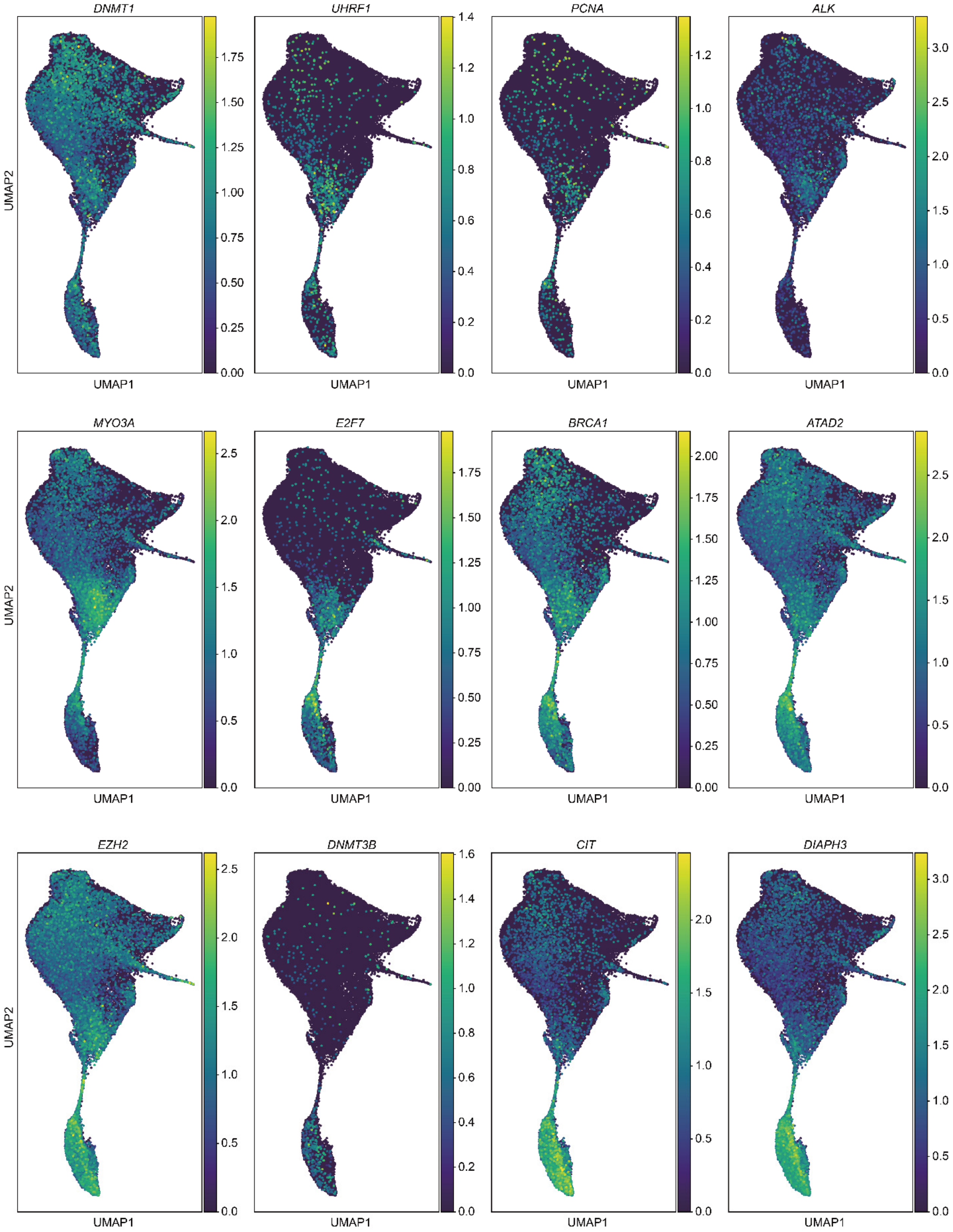


**Figure S16. Gene expression profiles associated with path 1 of the tumor population in GIST.**

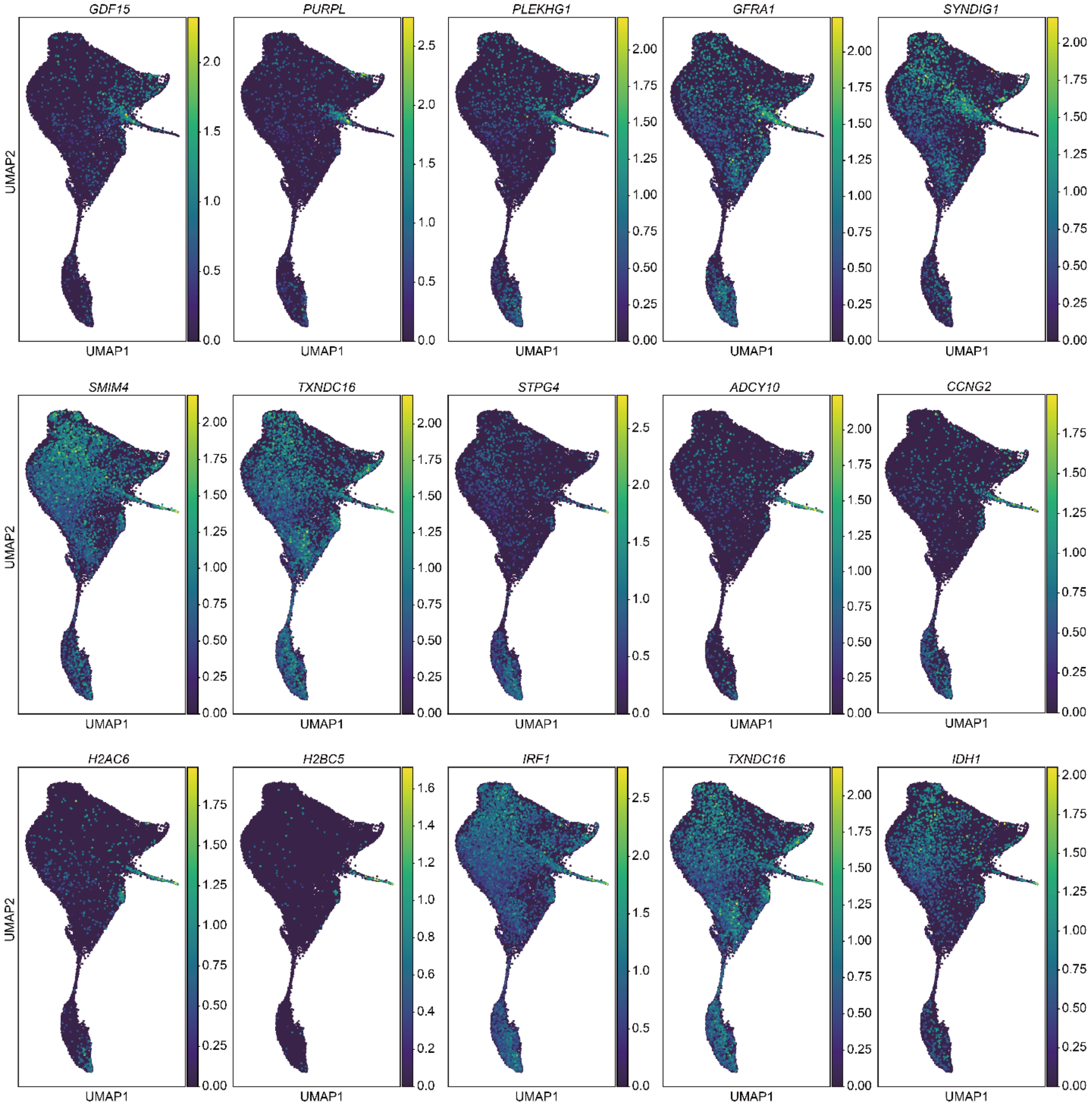


**Figure S17. Gene expression profiles associated with path 2 of the tumor population in GIST.**

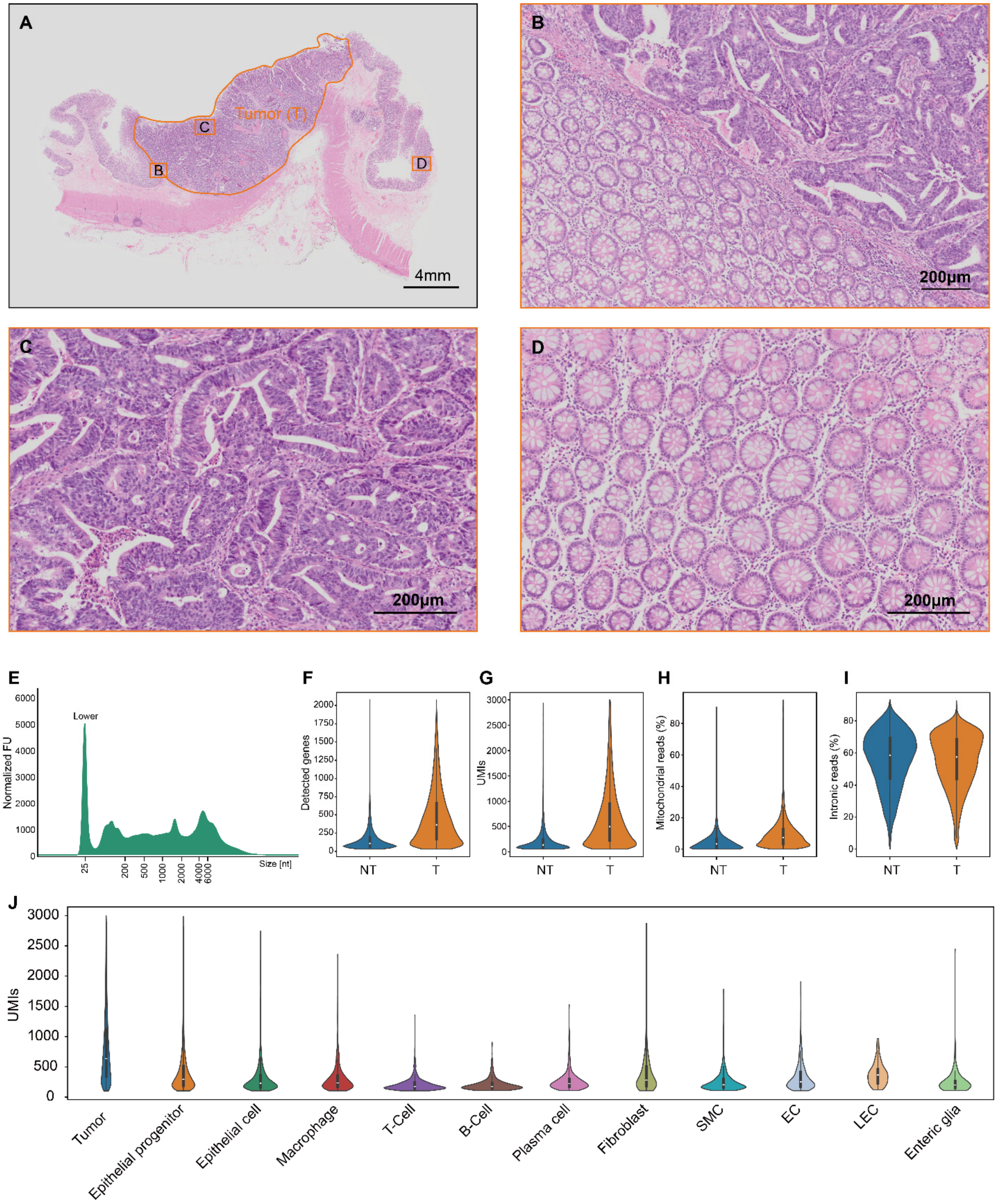


**Figure S18. Analysis result of FX-seq in archived CRC FFPE tissues.**

(A) Overall microscopic image of the H&E-stained section of CRC FFPE.

(B–D) Magnified images of different morphologic regions. (B) shows the border between non-tumor (NT) and tumor (T) region. (C) shows tumor regions with collapsed mucosal tissue morphology, whereas (D) shows non-tumor regions with intact tissue morphology.

(E) Automated gel electrophoresis result of total RNA extracted from the FFPE section of the CRC specimen. Data are plotted by normalized fluorescence units (FU) per size (bp). “Lower” indicates low-molecular-weight species markers.

(F–I) Quality metrics of the CRC FX-seq experiment comparing spatially resolved tumor and non-tumor regions. Detected genes (F), UMIs (G), fraction of mitochondrial reads (H), and fraction of intron-aligned counts (I).

(J) UMIs of annotated cell types in Figure 5.

Abbreviations are as follows: SMC, smooth muscle cell; EC, endothelial cell; LEC, lymphatic endothelial cell; Enteric glia, enteric glial cell.

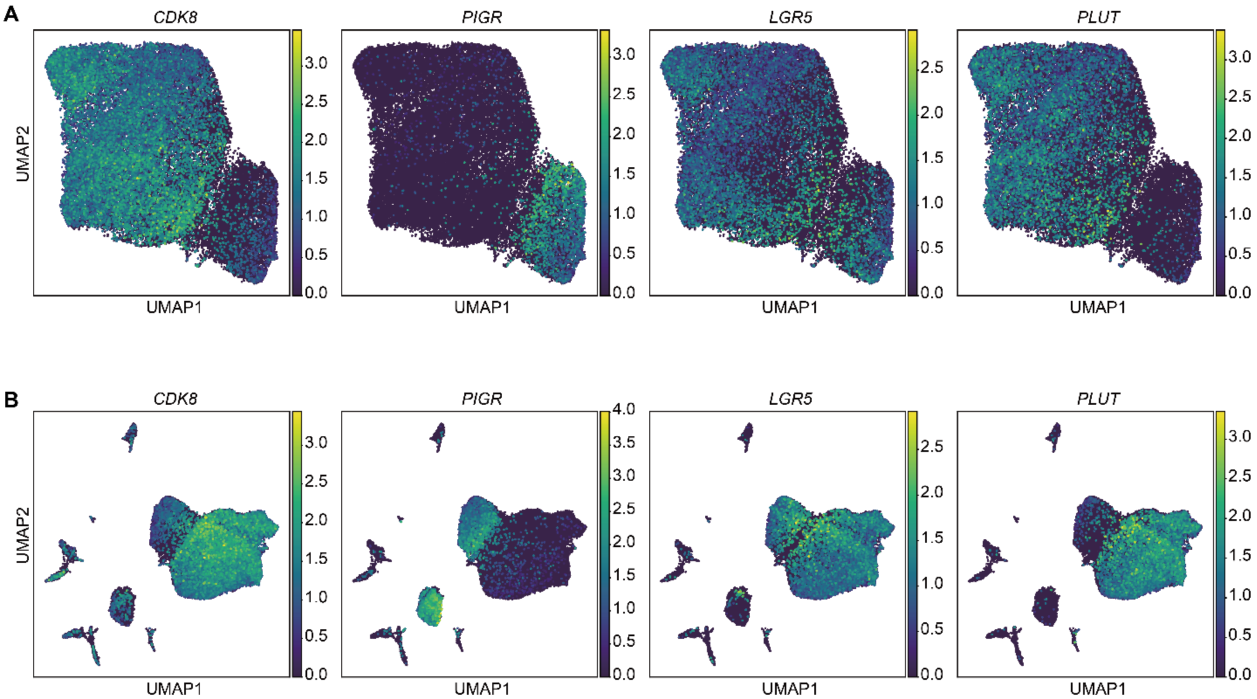


**Figure S19. Differential gene expression profiles of tumor populations in colorectal cancer (CRC).**

(A) Gene expression in subclusters, including epithelial progenitor and tumor populations.

(B) Gene expression in all identified populations.

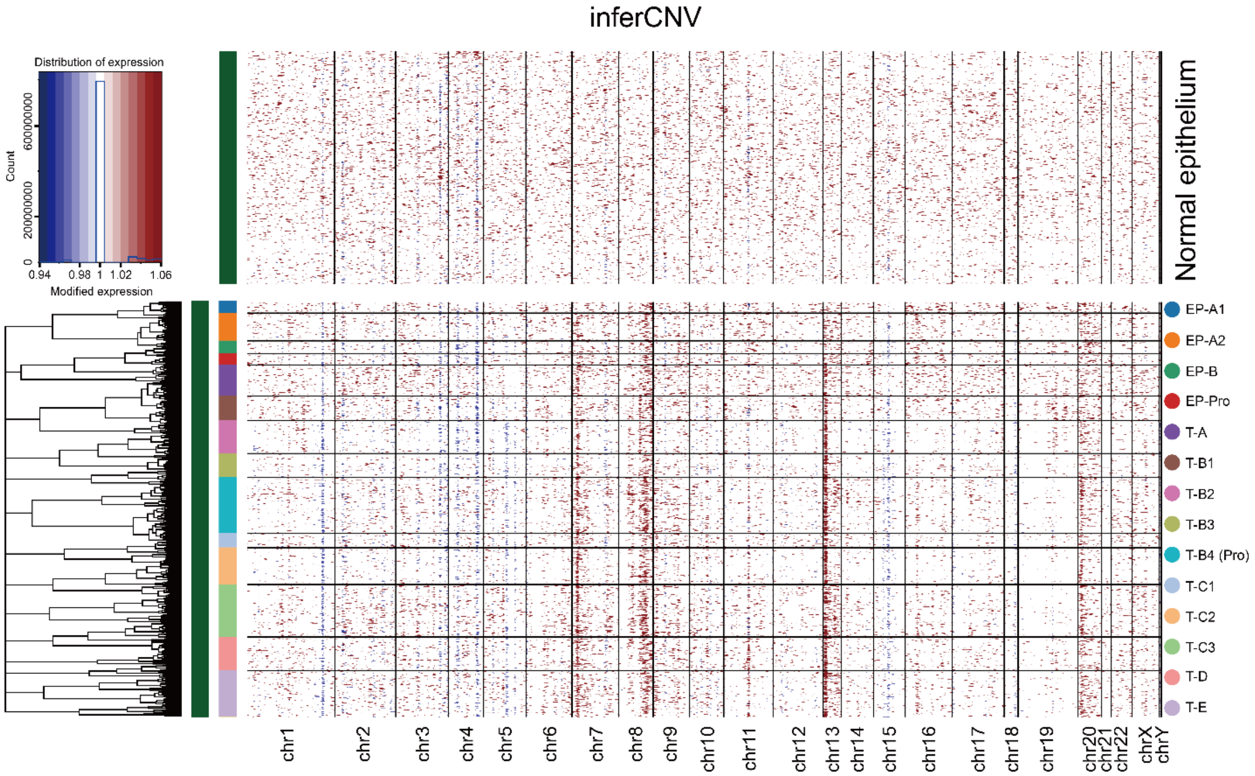


**Figure S20. InferCNV analysis result of individual cells.**

Legends for Tables S1-17

**Table S1. Differentially expressed genes across annotated cell types in the no treatment control group of PFA-perfused mouse brain related to Figure 1.**

**Table S2. Differentially expressed genes across annotated cell types of FX-seq group of PFA perfused mouse brain related to Figure 1.**

**Table S3. Differentially expressed genes across annotated cell types from freshly extracted and lightly fixed cerebellum removed mouse brain related to Figure 2.**

**Table S4. Differentially expressed genes across annotated cell types from FX-seq of heavily fixed cerebellum removed mouse brain related to Figure 2.**

**Table S5. Differentially expressed genes across annotated cell types from no treatment control of mouse brain FFPE block related to Figure S10.**

**Table S6. Differentially expressed genes across annotated cell types from FX-seq group of mouse brain FFPE block related to Figure S10.**

**Table S7. Differentially expressed genes across annotated cell types from mouse brain FFPE section related to Figure 3.**

**Table S8. Differentially expressed genes across annotated cell types from mouse brain H&E section related to Figure 3.**

**Table S9. Differentially expressed genes across annotated cell types from FFPE and H&E sections of omental metastasis from bladder cancer related to Figure 3.**

**Table S10. Differentially expressed genes across annotated cell types from the gastrointestinal stromal tumor (GIST) FFPE block related to Figure 4.**

**Table S11. Differentially expressed genes across leiden clusters from subcluster analysis of tumor population of GIST related to Figure 4.**

**Table S12. Differentially expressed genes across annotated cell types from FFPE sections from colorectal cancer (CRC) related to Figure 5.**

**Table S13. Differentially expressed genes across annotated cell types from subcluster analysis of fibroblast population of CRC related to Figure 5.**

**Table S14. Differentially expressed genes across annotated cell types from subcluster analysis of tumor population of CRC related to Figure 5.**

**Table S15. Total reads, detected cell number, and reads per cell from each sequencing result.**

**Table S16. Scanpy function parameters used for the analytical pipeline.**

**Table S17. List of oligonucleotide sequences.**
